# A single wave of monocytes is sufficient to replenish the long-term Langerhans cell network after immune injury

**DOI:** 10.1101/617514

**Authors:** Ivana R. Ferrer, Heather C. West, Stephen Henderson, Dmitry S. Ushakov, Pedro Santos e Sousa, Jessica Strid, Ronjon Chakraverty, Andrew J. Yates, Clare L. Bennett

**Affiliations:** Institute of Immunity and Transplantation, Division of Infection and Immunity, University College London, London NW3 2PF, UK; Haematology, Division of Cancer Studies, University College London, London WC1E 6DD, UK; Peter Gorer Department of Immunobiology, School of Immunology and Microbial Sciences, King’s College London, New Hunt’s House, Newcomen St, London SE1 1UL, UK; Division of Immunology and Inflammation, Imperial College London, Hammersmith campus, London W12 0NN, UK; Department of Pathology and Cell Biology, Columbia University Medical Center, New York, NY10032, USA

**Keywords:** Langerhans cells, monocytes, epidermis, skin, bone marrow transplant

## Abstract

Embryo-derived Langerhans cells (eLC) are maintained within the sealed epidermis without contribution from circulating cells. When the network is perturbed by transient exposure to ultra-violet light, short-term LC are temporarily reconstituted from an initial wave of monocytes, but thought to be superseded by more permanent repopulation with undefined LC precursors. However, the extent to which this mechanism is relevant to immune-pathological processes that damage LC population integrity is not known. Using a model of allogeneic hematopoietic stem cell transplantation, where allo-reactive T cells directly target eLC, we have asked if and how the original LC network is ultimately restored. We find that donor monocytes, but not dendritic cells, are the precursors of the long-term LC in this context. Destruction of eLC leads to recruitment of a single wave of monocytes which engraft in the epidermis and undergo a sequential pathway of differentiation via transcriptionally distinct EpCAM^+^ precursors. Monocyte-derived LC acquire the capacity of self-renewal, and turn-over in the epidermis was remarkably similar to that of steady state eLC. However, we have identified a bottleneck in the differentiation and survival of epidermal monocytes, which together with the slow turn-over of mature LC limits repair of the network. Furthermore, replenishment of the LC network leads to constitutive entry of cells into the epidermal compartment. Thus, immune injury triggers functional adaptation of mechanisms used to maintain tissue-resident macrophages at other sites, but this process is highly inefficient in the skin.

**Highlights:**

- Immune injury leads to recruitment of a single wave of monocytes to replace resident Langerhans cells (LC).
- DC lineage cells cannot become long-term replacement LC.
- The size of the re-emerging network is controlled by density-dependent division of mature LC.
- Immune injury and inefficient repopulation by monocyte-derived cells lead to a permanently altered LC niche.

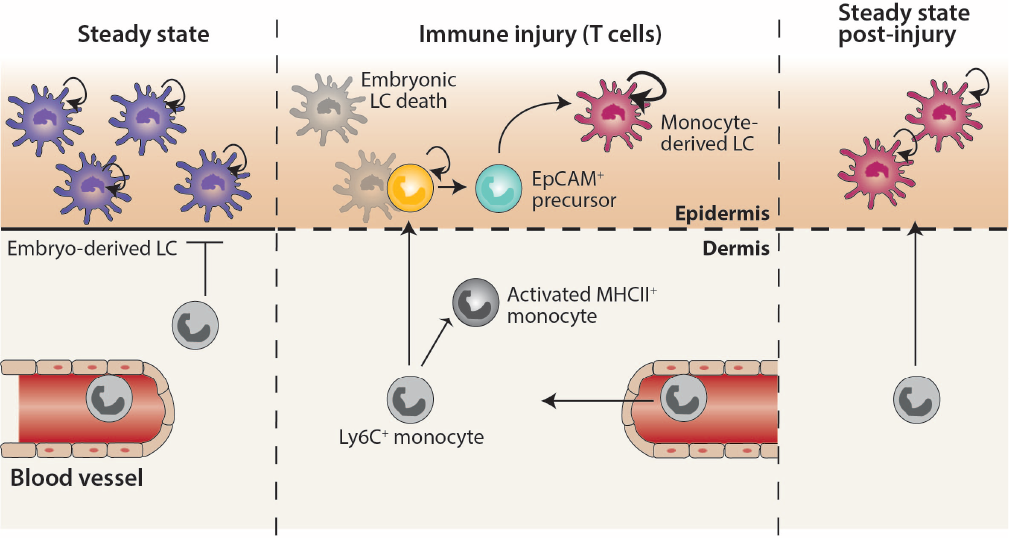

## Introduction

Langerhans cells (LC) are unique mononuclear phagocytes that reside within the epithelial layer of the skin and mucosal tissues (1, 2), where they play a key role in regulating immunity at the barrier surface. Loss of LC-dependent immune surveillance leads to a break-down in skin tolerance, and increased susceptibility to infection (3). Within the skin, embryonic (e)LC differentiate from yolk sac and fetal common tissue macrophage precursors that seed the skin before birth (4–8). The density of the adult eLC network is established post-birth by a burst of local proliferation of differentiated LC in response to undefined signals (9). Subsequently, the mature network is maintained by low levels of clonal cell division within adult skin (10–12). By contrast, oral mucosal epithelia are populated by LC-like cells that are continuously seeded from recruited monocytes and dendritic cell (DC) precursors (1). These cells transcriptionally and phenotypically resemble epidermal eLC, but show little evidence for proliferation *in situ*, and rather depend on recruitment of blood-derived cells to maintain the cellular niche (1, 13).

Ablation of eLC from genetically engineered mice leads to patchy repopulation of the epidermis (14), due to division of surviving eLC with some contribution from bone marrow (BM) cells (12, 15). In this non-inflamed context, the replenishment of the empty niche is characterized by the slow kinetics by which emerging LC expand to fill the epidermis, reflecting the quiescent nature of mature LC within the skin environment (3). By contrast, severe perturbation of eLC in the context of inflammation leads to recruitment of BM-derived cells into the epidermis, and LC repopulation (10, 16). The cellular mechanisms by which this occurs have largely been defined using models in which transient acute exposure of murine skin to UV irradiation leads to cell death within the epidermis and eLC replacement. Under these conditions, Gr-1^+^ monocytes are recruited to the epidermis (17, 18). However, while some studies suggest that these monocytes can differentiate into long-term LC (17, 19), others have proposed a two wave model of eLC replacement, in which monocytes can persist for up to 3 weeks as ‘short-term’ LC-like cells, but are superseded by undefined precursors which become long-lived replacement LC (18). Central to this model is the observation that LC require the transcription factor Id2 for their development and persistence as long-lived quiescent cells (18). However, it remains controversial whether Id2 is required for repopulation of the LC niche after UV-irradiation (20). Furthermore, the nature of the long-term LC precursors remains a key question in the field, with a number of studies suggesting the potential for dendritic cells (DC) or their precursors, to seed epidermal LC in the adult (21–25).

The resident macrophage population in most tissues is maintained by recruitment of Ly6C^+^ classical monocytes from the blood (26). Ly6C^+^ monocytes are short-lived, non-dividing cells once they have left the BM; however they show remarkable plasticity upon differentiation within tissues (27, 28). One active area of investigation is whether monocyte-derived macrophages can transcriptionally and functionally replace the resident macrophage populations that were originally seeded from intrinsically distinct precursors at birth. In the steady state, genetic ablation of tissue resident cells results in the differentiation of Ly6C^+^ monocytes into Kupffer cells (29) and alveolar macrophages (30) in the liver and lung, respectively, that show few differences from the cells they replace. By contrast, genetic ablation of microglia leads to repopulation by monocyte-derived cells that appear to fulfil the functional roles of their resident counterparts, but remain morphologically distinct, and continue to express monocyte-related genes (31, 32). Whether re-emerging skin LC, driven by immune injury and inflammation in the skin, also retain evidence of their cellular origin, and to what extent repopulating cells can become a long-term quiescent LC network, remain pertinent questions. Not least because monocyte-derived cells in other inflamed tissues remain transcriptionally and functionally distinct from resident cells (33, 34).

We have exploited a murine model of hematopoietic stem cell transplant to define the cellular mechanisms that control re-building of the LC network after immune-mediated pathology. We demonstrate that T cell-mediated destruction of eLC leads to the recruitment of a single wave of monocytes into the epidermis. Here, monocytes proliferate and differentiate into unique EpCAM^+^ cells that seed long-term monocyte-derived (m)LC, which are indistinguishable from the embryo-derived cells that have been replaced. We provide evidence for a surge of monocyte recruitment to the epidermis, but differentiation of these cells into mLC is a significant rate limiting step, resulting in inefficient rebuilding of the mature LC network. In addition, the epidermal compartment is not re-sealed after entry of T cells, and remains open to circulating cells. Thus, immune injury triggers an adaptive process that converges with mechanisms to regenerate other tissue-resident macrophages, but this is highly inefficient in the skin.

## Results

### Immune injury leads to the gradual replenishment of the epidermis with LC-like cells

Epidermal eLC are destroyed within a week of exposure to UV irradiation, and the niche is fully repopulated by BM-derived cells 2 weeks later (10). To compare the kinetics of LC turn-over after immune injury, we used a murine model of minor histocompatibility antigen mis-matched BM transplantation (BMT) (Figure S1). eLC are radio-resistant (16) and persisted in control mice that received BMT alone. A small population of donor BM-derived cells was evident in some mice 10 weeks post-transplant, but we observed significant variability between mice (Figure 1A top panel and B). By contrast, transfer of CD8^+^ male antigen-reactive Matahari (Mh) T cells with BMT (35, 36) led to the entry of pathogenic T cells into the skin epidermis, and destruction of host male eLC (37). Loss of eLC was complimented by the emergence of donor BM-derived CD11b^+^Langerin^+^ LC-like cells in the epidermis 2 weeks post-transplant, which increased sharply in frequency and number between 2-3 weeks (Figure 1A bottom panel and B), concomitant with the peak of Mh T cell numbers in the epidermis (Figure 1C). The development of full donor LC chimerism was gradual and evident by week 10 post-transplant. At this time-point repopulating donor CD11b^+^Langerin^+^ LC-like cells were phenotypically indistinguishable from host eLC in BMT controls according to markers defined in previous studies (Figure 1D) (18, 38).

**Fig. 1.**
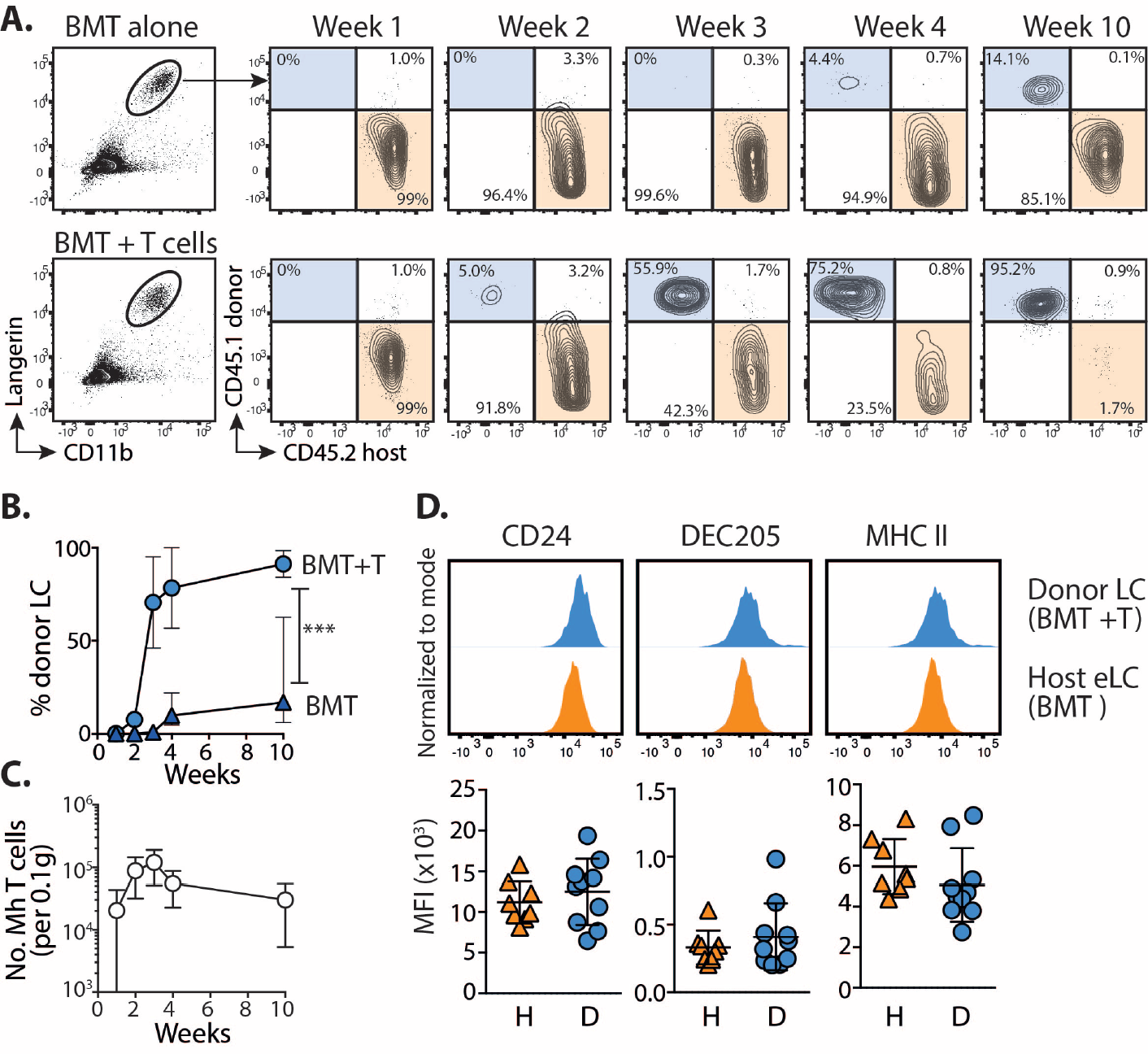
Immune injury leads the gradual replenishment of the epidermis with LC-like cells. **A**. Male recipients received female bone marrow alone (BMT) or with CD4 and CD8 (Matahari) T cells. Chimerism was measured within the mature CD11b^+^Langerin^+^ LC population at different time points. Representative flow plots show the relative frequency of host (CD45.2) and donor (CD45.1)-derived cells. **B**. Graph showing the frequency ± SD of donor LC in mice receiving BMT with (circles) or without (triangles) T cells. 2-way ANOVA P<.0001. Data are pooled from 2 independent experiments for each time point (n=5-10). **C**. Graph shows the number ± SD of Vβ8.3^+^ Matahari (Mh) T cells in the epidermis over time, per 0.1g total ear tissue (n=7-8). **D**. Top – representative histogram overlays show the expression of LC-associated proteins on donor-derived LC (from mice that received BMT + T cells) or host eLC (BMT alone), 10 weeks post-transplant. Bottom – summary data showing the median fluorescent intensity (MFI) for each sample. H = host, D = donor, each symbol is one mouse. Data are pooled from 2 independent experiments (n=8), and representative of >3 different experiments.

Therefore, infiltration of Mh T cells into the epidermis leads to the destruction of resident host eLC and gradual repopulation by donor BM-derived LC.

### DC lineage cells do not become long-term replacement LC

The DC-like nature of LC has led to the suggestion that DC lineage cells may contribute to adult LC, and/or seed repopulating cells after damage (21, 23). This hypothesis has recently been supported by work demonstrating that circulating human CD1c^+^ DC have the potential to become LC-like cells *in vitro*, but this has not been directly tested *in vivo* (24, 25). Therefore, we investigated the possibility that DC lineage cells may contribute to LC repopulation after immune injury, by transplanting irradiated male recipients with a 1:1 mixture of BM from Vav-Cre.Rosa26LSL^Tomato^ (Vav^Tom^) and Clec9a-Cre.Rosa26LSL^EYFP^ (Clec9a^YFP^) reporter lines. The Vav-Cre transgene is expressed by all hematopoietic cells (39) and provides an internal control for the development of BM-derived LC, while Clec9a-dependent YFP marks cells restricted to the DC lineage (40) (Figure 2A). 10 weeks later, approximately 60% of splenic CD11c^+^MHCII^+^ cells were derived from Vav^Tom^ BM and all expressed Tomato, while about half of the Clec9a^YFP^-derived cells expressed YFP. By contrast, while there was a clear contribution of donor Vav^Tom^ cells repopulating LC after BMT with T cells, we did not detect any YFP^+^ LC (Figures 2B and C).

**Fig. 2.**
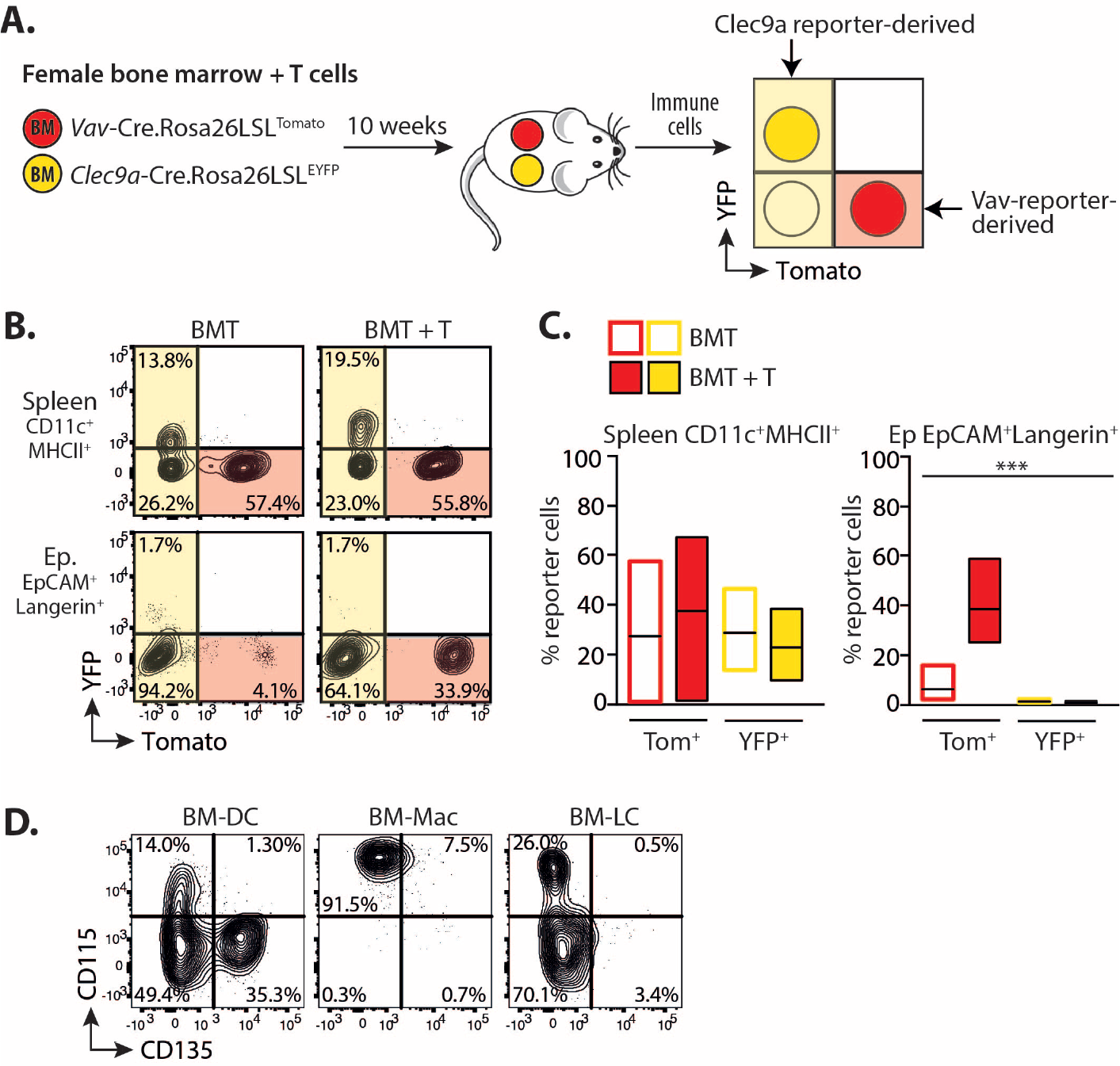
DC lineage cells do not become long-term replacement LC. **A**. Schematic showing the experimental procedure. Male mice received female BMT with T cells. BM was composed of a 1:1 mixture of cells from syngeneic female Clec9a^YFP^ and Vav-^Tom^ mice. 10 weeks later splenocytes and epidermal LC were assessed for the relative contribution of cells expressing Tomato (Tom) or YFP. **B**. Representative contour plots showing gated CD11c^+^MHCII^+^ cells in the spleen or CD11b^+^EpCAM^+^Langerin^+^ LC in the epidermis of mice that received BMT with or without T cells. **C**. Summary bar graphs showing the frequency of red Tom^+^ or yellow YFP^+^ cells within splenic CD11c^+^MHCII^+^ (left) or epidermal EpCAM^+^Langerin^+^ (right) cells in mice receiving BMT alone (open bars) or BMT with T cells (filled bars). Bars show the mean and range of data points. Data are pooled from 2 independent experiments, and analysed using a 2-way ANOVA P<.0001 (n = 5-6). **D**. BM cells were cultured with either GM-CSF to generate BM-DC (MHCII^high^CD11c^+^DEC205^low to +^EpCAM^neg^) and macrophages (MHCII^int^CD11c^+^DEC205^low to +^EpCAM^neg^ CD11b^high^) or GM-CSF and TGFβ to generate BM-LC (MHCII^int^CD11c^+^ DEC205^+^EpCAM^+^CD11b^low^). Representative contour plots show the percentage of CD135- and CD115-expressing cells in the gated populations. Data are representative of 1 experiment with BM-DC and Macs, and 3 independent experiment for BM-LC on days 4-6.

The ontogeny of DC lineage cells and monocyte-derived (mo)DC can be delineated in BM cultures by tracking residual expression of the DC and monocyte/macrophage-defining receptors fms-like tyrosine kinase (Flt)-3 and CSF1R (CD115) respectively (41, 42). Culture of BM cells with GM-CSF led to the expansion of Flt3^+^ and CD115^+^ cDC and moDC, and CD115^+^ macrophages, based on the gating strategy published by the Reis e Sousa group (Figure S2) (41). By comparison, culture of BM cells with GM-CSF and TGFβ to generate BM-LC (20, 43, 44) demonstrate a more heterogeneous profile within EpCAM^+^DEC205^+^ cells with subsets of both CD115^+^ and CD115^neg^ cells. However, there was no contribution of CD135^+^ cells to the LC-like cells in these cultures, mirroring our *in vivo* findings (Figure 2D).

Therefore, in conclusion we find no evidence for the contribution of DC lineage cells to BM-derived LC *in vitro*, or the long-term replacement LC network *in vivo*.

### LC repopulation is preceded by a single wave of donor CD11b^+^ cells

Mature LC are identified by the unique concomitant expression of high levels of the cell adhesion molecule EpCAM (CD326) and the C-type lectin receptor, Langerin (CD207), which are simultaneously up-regulated upon differentiation of eLC (9, 19, 43). By contrast, ‘short-term’ LC, do not up-regulate EpCAM (18). We observed the T cell-dependent accumulation of CD11b^int to high^ cells within the epidermis (Figures 3A and B), that peaked 3 weeks post-transplant and preceded the shift to donor LC chimerism shown in Figure 1A. Phenotypic analysis of these cells demonstrated the presence of 3 populations sub-divided by expression of EpCAM and Langerin (Figure 3C): donor CD11b^high^ cells contained 2 sub-populations that were either negative for both markers, suggesting recent arrival in the epidermis, or solely expressed EpCAM; by comparison CD11b^int^ cells were EpCAM^high^Langerin^high^ and therefore resembled mature LC. For this paper, we will refer to these populations as ‘CD11b^high^’, ‘EpCAM^+^’ and ‘donor LC’, respectively. We observed a dramatic peak in the frequency and number of EpCAM^+^ cells between 2-3 weeks post-transplant. These data strongly suggested a developmental trajectory whereby a single wave of epidermal CD11b^high^ cells was sufficient to repopulate the mature LC network. We therefore reasoned that acquisition of LC-defining proteins would be consistent with the developmental transition of these sub-populations. Indeed, we observed the gradual loss of CD11b, and up-regulation of CD24 and DEC205 with differentiation of donor LC (Figure 3D).

**Fig. 3.**
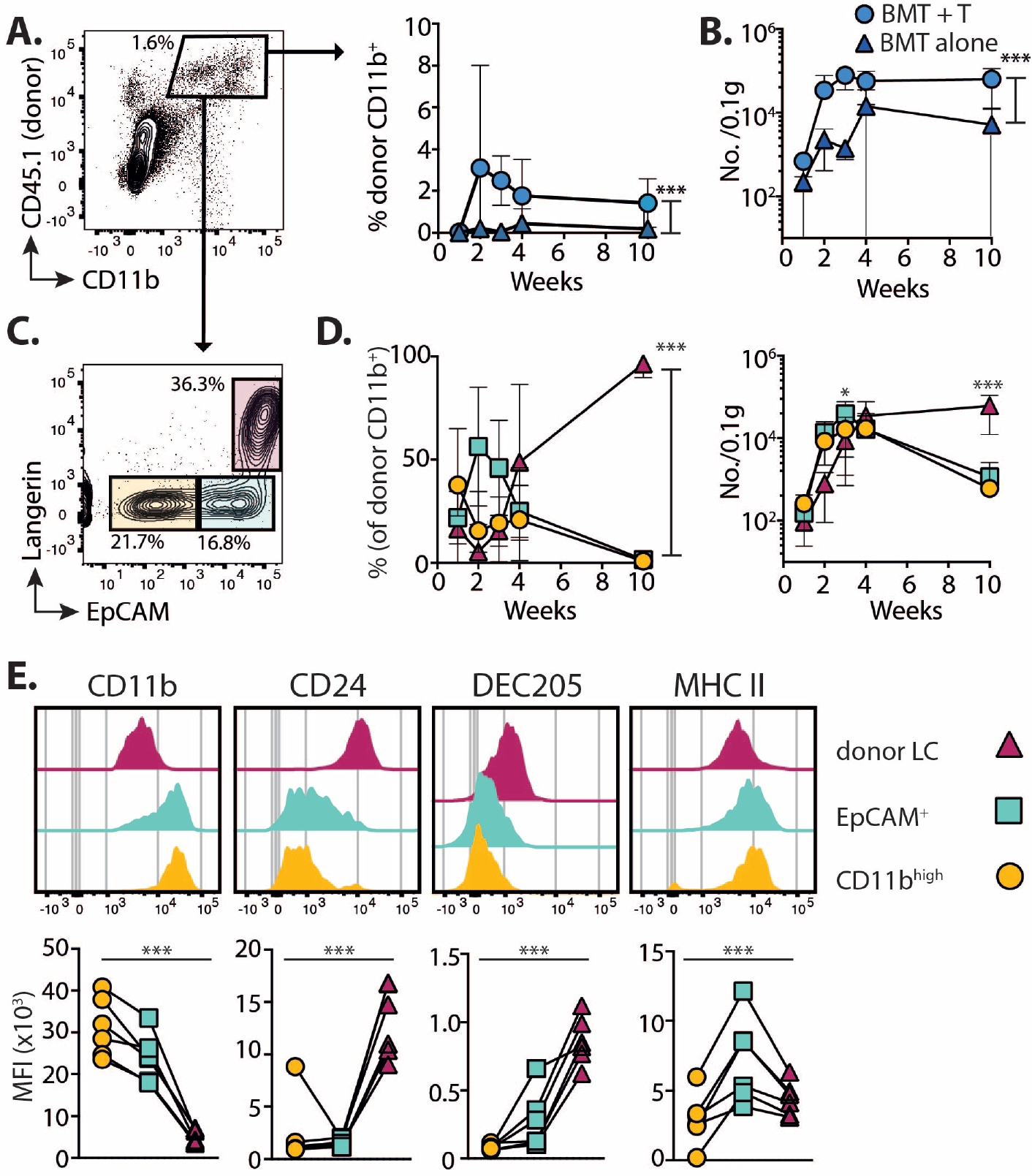
LC repopulation is preceded by a single wave of CD11b^+^ cells. Mice received BMT with T cells, and the epidermis was analysed at different time points. **A**. Left – dot plot shows the gating of single CD11b^int to high^CD45.1^+^ donor myeloid cells. Right – summary graph showing the frequency ± SD donor CD11b^+^ cells in mice receiving BMT alone (triangles) or BMT + T cells (circles). P=.0004 (n= 6-7). **B**. Graph shows the number ± SD per 0.1g total ear weight of donor CD11b^+^ cells, P<.0001 (n=5-10). **C**. Representative contour plots at 3 weeks showing 3 distinct sub-populations within single CD11b^int to high^CD45.1^+^ cells. **D**. Summary graphs showing the frequency ± SD (left) and number ±SD (right) of cells within each of the gated populations shown in C: Circles CD11b^high^ (EpCAM^neg^Langerin^neg^); squares EpCAM^+^; triangles donor LC (EpCAM^+^Langerin^+^). (n=7-8, pooled from 2 independent experiments and representative of >5 different experiments). P<.0001 for frequency 2-way ANOVA; significance for numbers was calculated with a 2-way ANOVA with Tukeys multiple comparisons test; at 3 weeks P=.0196 and at 10 weeks P<.0001. **E**. Top – representative histogram overlays show surface expression levels of LC-defining proteins in the gated donor populations 3 weeks post-transplant. Bottom – graphs show summary data for the median fluorescent intensity (MFI). Symbols represent individual samples, P<.0001, or. 0136 for MHCII with a 1-way ANOVA. Data are pooled from 2 independent experiments per time point (n=6).

Together, our data demonstrate that re-building of the LC network is solely dependent on a wave of CD11b^+^ myeloid cells that differentiate *in situ* to become LC. The disconnect between EpCAM and Langerin expression enabled us to subsequently define the process by which donor LC emerged from EpCAM^+^ cells in more detail.

### EpCAM^+^ monocyte-derived cells are distinct from donor LC

Our phenotyping data suggested that incoming CD11b^+^ cells differentiated into a unique EpCAM^+^ intermediate before becoming donor LC, but it was possible that EpCAM^+^ cells were already immature LC. To distinguish between these possibilities, we compared the transcriptional profile of EpCAM^+^ cells to donor LC or other CD11b^+^ populations in the skin and blood (Figure 4A and see Figure S3 for sorting strategy). Hierarchical clustering demonstrated that eLC and donor LC were interchangeable, but that EpCAM^+^ cells clustered as a distinct population, and were more closely aligned to blood and dermal monocytes than mature LC (Figure 4B). Analysis of the genes that contributed to differences along the PC1 axis after principle components analysis (Figure S4A) demonstrated that EpCAM^+^ cells were distinguished by the down-regulation, but not loss, of expression of genes associated with monocyte development and function (e.g. *Prr5*, *Ccl9*, *Fcgr3*, *Fcgr4*, *Trem3*, *Tlr7*) (Figure S4B), including *Trem14*, which is expressed by Ly6C^+^ monocytes with the potential to become moDC (45). However, EpCAM^+^ cells had not up-regulated genes associated with changes to cell structure, adhesion and signalling that defined donor LC (e.g. *Emp2*, *Kremen2*, *Nedd4*, *Ptk7*). Given the distinct clustering of monocytes/EpCAM^+^ cells and eLC/donor LC, we directly tested the contribution of monocytes to emerging LC by transferring mixed congenic Ccr2^+/+^ and syngeneic Ccr2^−/−^ BM with T cells (Figure 4C). CCR2 is required for both egress of monocytes from the BM and entry into tissues (46). Our data clearly showed that only Ccr2^+/+^ cells contributed to CD11b^high^, EpCAM^+^ and donor LC sub-populations (Figures 4D and E), demonstrating a monocytic origin for these cells.

**Fig. 4.**
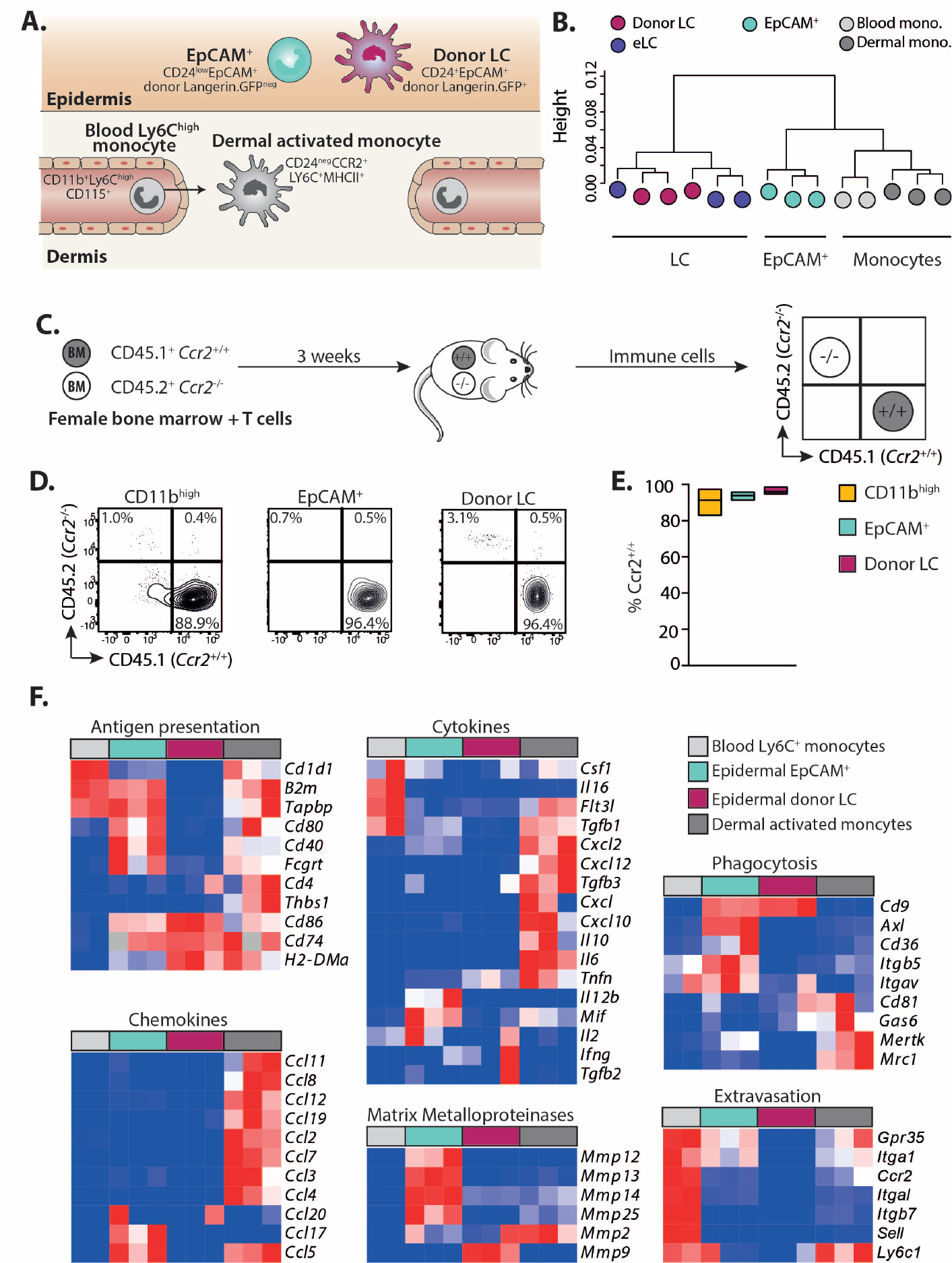
EpCAM^+^ monocyte-derived cells are distinct from donor LC. **A**. Schematic showing the populations of cells and phenotypic markers used to isolate cells for sequencing. **B**. Dendrogram showing clustering of samples. **C**. Schematic illustrating competitive chimera experiments to test the requirement for monocyte-derived cells. Male mice received female BMT with T cells. BM was composed of a 1:1 mixture of cells from congenic wild-type (CD45.1^+^*Ccr2*^+/+^) or CCR2-deficient (CD45.2^+^*Ccr2*^−/−^) mice. Epidermal cells were analysed 3 weeks later. **D**. Representative contour plots showing the frequency of wild-type or knock-out cells within gated donor epidermal myeloid cells (host cells were excluded at this time point by the use of Langerin.EGFP recipients, and exclusion of GFP^+^ LC from our analyses). **E**. Summary data showing the frequency of CCR2^+/+^, donor cells within each population. Bar graphs show the mean and range of data points, data are pooled from 2 independent experiments (n=6). Percent of *Ccr2*^+/+^ cells versus *Ccr2*^−/−^ in each population P<.0001, One-way ANOVA. **F**. Heat maps showing relative gene expression of defined genes grouped into panels according to distinct functional processes. Blood monocytes (grey) n = 2, EpCAM^+^ cells (cyan) n = 3, donor LC (magenta) n = 3, dermal activated monocytes (grey) n = 3.

Ly6C^+^ monocytes mature into MHCII^+^ activated monocytes (or monocyte-derived DC) in the dermis (47). Therefore, we directly compared the outcomes of monocyte differentiation within different skin compartments using panels of genes associated with monocyte maturation and function described by Schridde and colleagues (48) (Figure 4F). This analysis demonstrated the divergence between Ly6C^+^ monocytes differentiating within the dermis or epidermis. EpCAM^+^ cells displayed a unique gene signature associated with tissue homeostasis and modulation of the epidermal niche by matrix metalloproteinases (*mmp12*, *13*, *14*, and *25*, but not *mmp2* and *9* which are associated with egress of mature LC out of the epidermis (49)); phagocytosis and uptake of apoptotic cells (*cd9*, *axl*, *cd36*, *itgb5*, *itgav*); and activation of complement (*clqa*, *clqb*, *clqc*). However, EpCAM^+^ also retained shared patterns of gene expression with moDC that suggested recent extravasation from the blood (*gpr35*, *itgal*, *ccr2*), and the potential to activate T cells (*b2m*, *tapbp*, *cd80*, *cd40*, *fcgrt*), which was lacking from LC.

Thus, monocytes replace eLC after T cell-mediated destruction of the network. Entry of monocytes into the epidermis triggers differentiation of a unique population of EpCAM^+^ cells that express a gene program which favors residency within the epidermis, before development into mature mLC.

### Proliferation of monocytes and mLC *in situ* combine to replenish the LC network

Our data implied that monocytes were sufficient to replenish the LC network, but Ly6C^+^ monocytes are short-lived non-cycling cells, while proliferation of eLC at birth and in adults determines the density LC within the epidermis (9, 12). Therefore, we considered the relative importance of recruitment versus proliferation of epidermal CD11b^+^ cells for the rebuilding of the LC network. We constructed mathematical models to quantify the flows between CD11b^high^ to EpCAM^+^ populations and the donor mLC pool using time courses of Ki67 expression (Figure S5). All epidermal populations showed evidence of active or recent cell division after transplant which decreased to homeostatic rates equivalent to eLC by 10 weeks (9.8 ± 1.49 percent (s.e.m.)(9). Models of the flow from CD11b^high^cells to EpCAM^+^ were fitted simultaneously to the time courses of total cell numbers and Ki67 expression in the two populations (Figure S5B). The kinetic of EpCAM^+^ cell numbers closely tracked that of the CD11b^high^ population, suggesting that EpCAM^+^ cells were short-lived and/or rapidly underwent onward differentiation (Table 1). We first explored whether the flow of CD11b^high^ and EpCAM^+^ cells into donor LC were consistent with a linear developmental pathway, as predicted by our experimental data. This was compared to the alternative scenario in which EpCAM^+^ cells were a ‘dead end’ population and wherein monocytes differentiated directly into LC, which we named the branched pathway (Figure 5A). The fitted predictions of the two models were visually similar (Figure S5C). Nevertheless, we found approximately 10-fold greater statistical support for the linear pathway, as measured by weights calculated using the Akaike Information Criterion (CITE) (50). Strikingly, however, maturation of EpCAM^+^ cells was highly inefficient with only 4% becoming donor mLC (Figure 5B and Table 1). Gene set enrichment analysis of EpCAM^+^ cells compared to donor LC suggested that most EpCAM^+^ cells underwent apoptosis within the epidermis (Figure S5D).

**Table 1.**
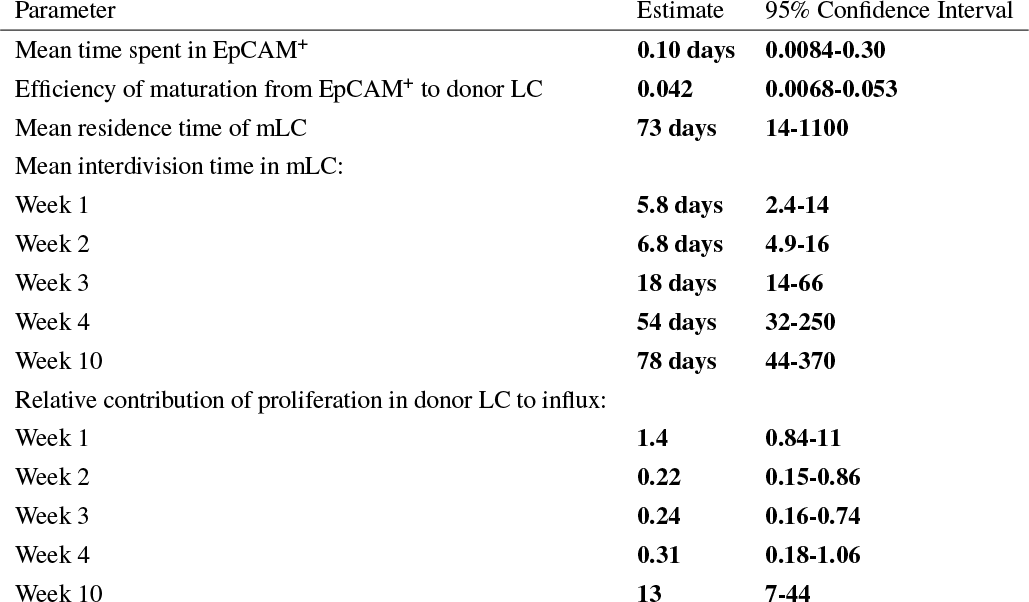
Parameter estimates from the best-fitting model describing the linear flow from incoming monocytes to CD11b^high^cells, EpCAM^+^ cells and to mature mLC.

| Parameter | Estimate | 95% Confidence Interval |
| --- | --- | --- |
| Mean time spent in EpCAM<sup>+</sup> | 0.10 days | 0.0084-0.30 |
| Efficiency of maturation from EpCAM<sup>+</sup> to donor LC | 0.042 | 0.0068-0.053 |
| Mean residence time of mLC | 73 days | 14-1100 |
| Mean interdivision time in mLC: |  |  |
| Week 1 | 5.8 days | 2.4-14 |
| Week 2 | 6.8 days | 4.9-16 |
| Week 3 | 18 days | 14-66 |
| Week 4 | 54 days | 32-250 |
| Week 10 | 78 days | 44-370 |
| Relative contribution of proliferation in donor LC to influx: |  |  |
| Week 1 | 1.4 | 0.84-11 |
| Week 2 | 0.22 | 0.15-0.86 |
| Week 3 | 0.24 | 0.16-0.74 |
| Week 4 | 0.31 | 0.18-1.06 |
| Week 10 | 13 | 7-44 |

**Fig. 5.**
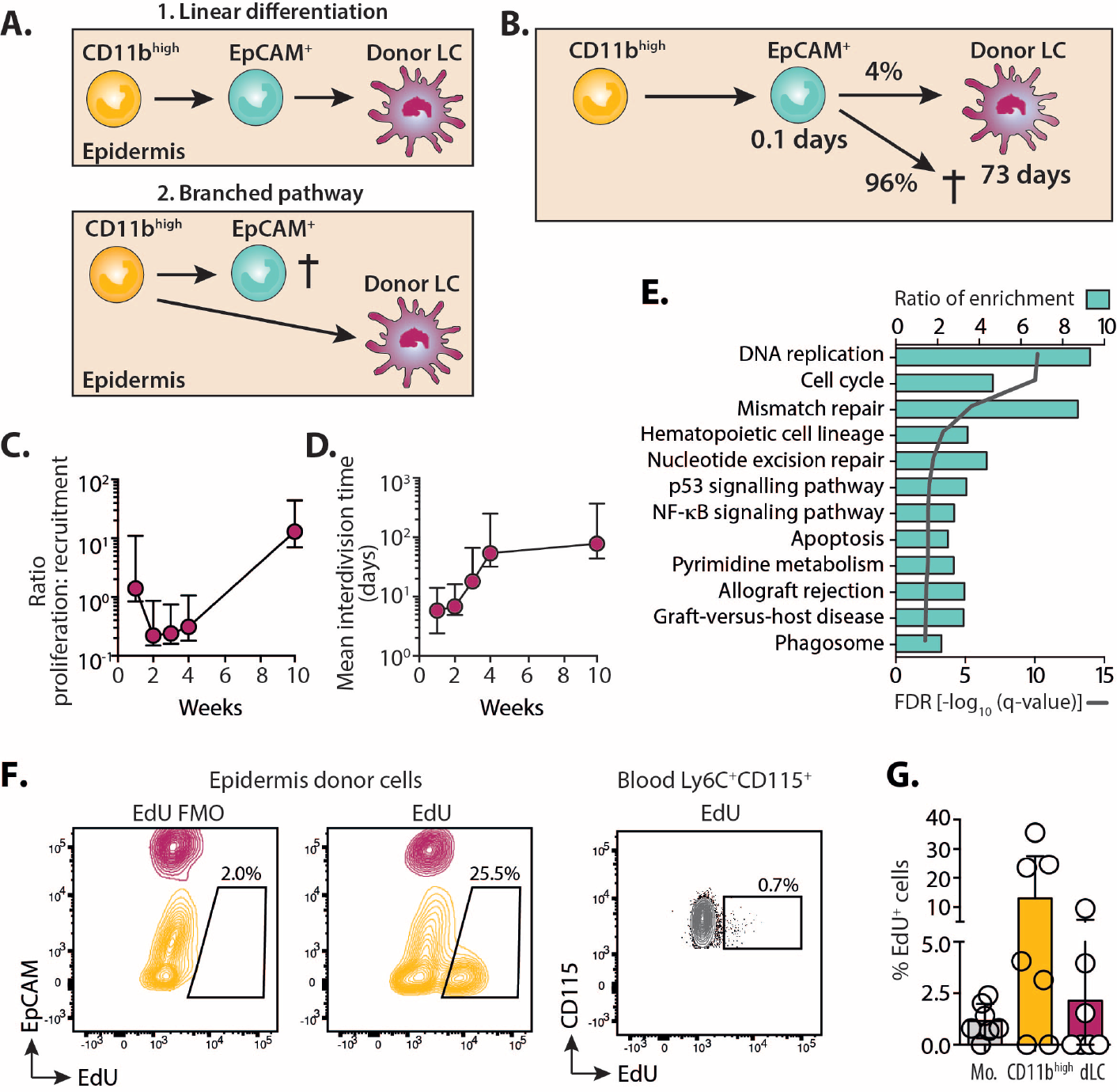
Proliferation of monocytes and LC *in situ* combine to replenish the LC network. **A**. Schematic to illustrate possible pathways of LC development from CD11b^high^ cells within the epidermis. **B**. Mathematical modelling fits strongly supported the linear development pathway. Numbers describe parameter estimates from the model. **C**. Graph showing the relative contribution of proliferation in donor LC to influx (± 95 percent confidence interval) over time. **D**. Graph showing the mean interdivisional time (± 95 percent confidence interval) of donor LC at different time points post-BMT with T cells. Parameter predictions from model fits are displayed in full in Table 1. **E**. Graphs showing the ratio of enrichment (bars) and FDR q values (line) for pathways predicted by WebGestalt to be over-expressed by EpCAM^+^ cells compared to blood monocytes. Overexpressed genes had a fold change more than or equal to 2, q < 0.05. **F**. Mice received EdU 3 weeks after BMT with T cells. 4 hours later, the skin and blood were harvested and cells analysed for incorporation of EdU. Representative contour plots show overlayed gated CD11b^high^Langerin^neg^ (yellow) or CD11b^+^Langerin^+^ (magenta) populations in the epidermis, or Ly6C^+^CD115^+^ monocytes in the blood. FMO is the fluorescent minus one stain without the EdU detection reagent. **G**. Summary graph showing the mean ± SD frequency of EdU^+^ cells in the different groups. Circles are individual mice, n=6. Data are pooled from 2 independent experiments. Mo. = monocytes, dLC = donor LC.

Production of donor LC was initially dominated by recruitment from CD11b^high^ and EpCAM^+^ cells but proliferative self-renewal replaced recruitment by week 10 when the mature LC pool had reached steady state (Figure 5C; Table 1). At this point, donor LC resided in the skin for approximately 10 weeks on average and divided once every 78 days (Table 1). These estimates were remarkably consistent with other observations of the rates of turnover and division of eLC in the steady state (10), and matched eLC doubling time within the unperturbed eLC network (12, 51). Fits were based on the assumption that the CD11b^+^ and EpCAM^+^ cells died or differentiated at a constant per cell rate. However, we had to include density-dependent proliferation of donor LC in order to explain the waning of Ki67^+^ cells over time. Thus, division occurred more frequently at low cell densities (Figure 5D; Table 1).

Expression of Ki67 by CD11b^high^ cells suggested that accumulation of LC precursors required local proliferation of undifferentiated cells. This hypothesis was supported by the over-representation of cell cycle pathways in EpCAM^+^cells (Figure 5E). This gene signature was in contrast to dermal monocytes, which up-regulated pathways associated with innate receptors and T cell activation (Figure S6). To pinpoint active cell division within epidermal cells *in vivo*, we injected EdU into mice 3 weeks after BMT with T cells, and analysed the frequency of EdU^+^ cells 4 hours later. Within this window, we detected incorporation of EdU by cycling CD11b^+^EpCAM^neg^cells in the epidermis, and less so in donor LC (Figures 5F and G).

Thus, modelling data strongly support the linear differentiation of monocytes in the epidermis. CD11b^high^ cells expand *in situ* to provide a large pool of LC precursors, but, differentiation of these cells into mLC is a bottleneck in the rate of replenishment of the LC network. Notably, we provide novel evidence of quorum sensing by mLC such that the rate of division slows as the niche is filled.

### Long-term mLC are homologous to eLC and up-regulate Id2

To determine how much of the eLC transcriptional profile was determined by origin we compared the transcriptional profiles of long-term (10 weeks) mLC to eLC from age-matched untreated mice in more detail. Correlation analysis demonstrated that mLC were virtually indistinguishable from eLC (Figure 6A), and the few differentiatlly expressed genes up-regulated in mLC were dominated by functions associated with cell adhesion and motility, suggesting the positioning and establishment of mLC within the epidermis (Table S1). To understand whether emergence of long-term mLC depended on restoration of a steady state environment in the epidermis, we also compared eLC and mLC 3 weeks post-transplant, at which point chimerism was incomplete and both populations shared the same inflammatory environment. We again found that mLC were homologous to eLC (Figure S7A), but that eLC showed evidence of prolonged exposure to the inflammatory environment due to conditioning and T cell damage (Figure S7B). Consistent with this, eLC isolated from the epidermis 3 weeks post-transplant primed CD8 T cells more efficiently than donor mLC from the same environment (Figure S7C).

**Fig. 6.**
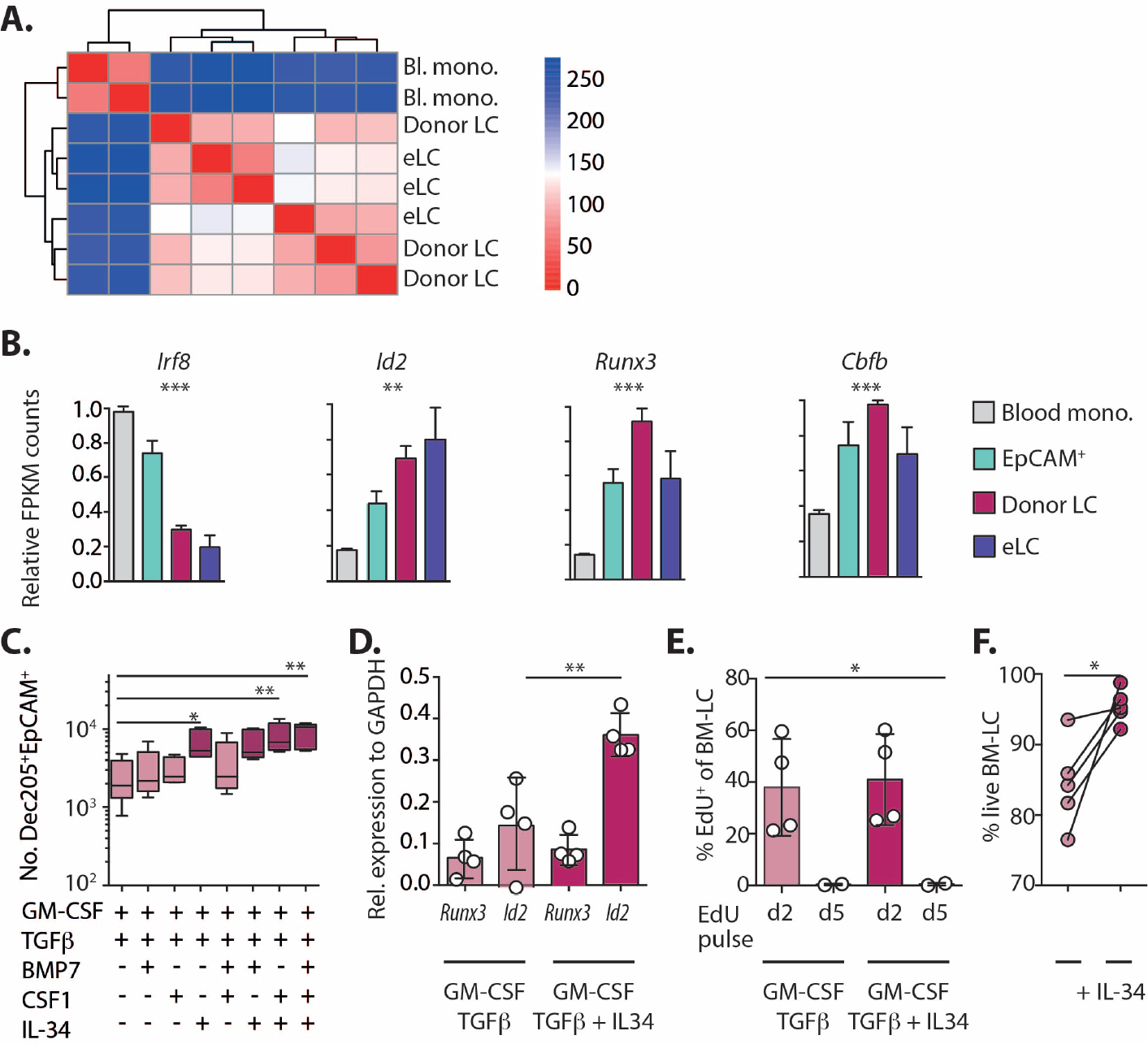
Long-term LC are homologous to eLC and up-regulate Id2. **A**. Correlation matrix comparing differentially expressed genes between and blood monocytes, mLC 10 weeks post-BMT + T cells, and eLC from age-matched controls. **B**. Graphs show the relative FPKM count normalised to the maximum value for different transcription factors from the RNAseq data. Significance was calculated with a 1-way ANOVA. *Irf8* P<.0001; *Id2* P=.0018; *Runx3* P=.0006; *Cbfb* P=.0032. Blood Ly6C^+^ monocytes n = 2, epidermal EpCAM^+^ cells n= 3, donor LC n = 3, age-matched eLC n = 3 **C**. BM cells were cultured for 6 days with GM-CSF, TFGβ and different combinations of BMP7, CSF1 and IL-34. Box and whiskers graph shows mean ± min. to max. numbers of DEC205^+^EpCAM^+^ cells in the cultures. GM-CSF + TGFβ alone versus addition of IL-34 P=.0469, versus IL-34 + CSF-1 P=.0013, versus IL-34, BMP7 and CSF-1 P=.0013. Significance was calculated with a 1-way ANOVA for non-parametric samples with Dunn’s multiple comparisons test. Each symbol is data from one culture, n = 5 independent BM donors, in 3 independent experiments. **D**. The bar graph shows the mean expression ± SD of *Runx3* or *Id2* relative to GAPDH in sorted DEC205^+^EpCAM^+^ cells. Symbols are cells from 4 independent BM donors, in 3 independent experiments. *Id2* expression in LC generated in the absence versus the presence of IL-34 P=.030, paired t test. **E**. Bar graph shows the mean frequency ± SD of EdU^+^ cells on day 6 of culture after cells where pulsed with EdU for 24 hours on day 2 or day 5. Symbols are cells from independent cultures (n = 2-4), 1-way ANOVA P=.024. **F**. Line graph shows the frequency of viable DEC205^+^EpCAM^+^ LC in GM-CSF / TGFβ cultures with, or without, IL-34. Symbols represent paired individual BM cultures, n = 5. Paired t-test, P=.0276.

Given that monocyte-derived cells rapidly differentiated into quiescent, long-lived LC, we reasoned that this must require programming by lineage-defining transcription factors (LDTF), namely Runx3 and Id2 in LC. Thus, we analysed expression these genes in mLC and their precursors. Figure 6B shows the sequential up-regulation of *Id2*, and *Runx3* and its partner *Cbf*β*2* (43), and down-regulation of monocyte-associated *Irf8*, as the cells became mLC. In addition, EpCAM^+^ cells showed evidence of early responsiveness to the dominant epidermal cytokine TGFβ (Figure S8A), and matured into LC-like cells upon culture with TGFβ *in vivo* (Figure S8B). Therefore, these data suggested that LDTF, and responsiveness to TGFβ, are switched on in EpCAM^+^ cells within the epidermal environment, before differentiation into mature LC. We next used an *in vitro* screen to identify the growth factors, in addition to TGFβ, that controlled Id2 expression and LC identity after differentiation from BM cells. We selected BMP7, CSF-1 and IL-34 based on expression of their cognate receptors (BMPR1a and CSF1R respectively) by EpCAM^+^ cells *in vivo* (Figure S8C) and their requirement for LC repopulation after UV-irradiation (43, 52, 53), and tested the impact of each factor on LC development in BM cultures. IL-34, but not CSF-1 or BMP7 specifically enhanced LC numbers (Figure 6C), and this was associated with the specific up-regulation of *Id2* by LC in IL-34 cultures (Figure 6D). BM-LC were derived from cells that proliferated before up-regulation of EpCAM in these cultures (Figure 6E), mirroring cycling of CD11b^high^ cells in the epidermis. However, cell division was not affected by addition of IL-34, which instead increased the survival of LC (Figures 6F).

Thus, monocytes differentiate into cells that express LDTF and become long-term LC that are homologous to eLC, with not evidence of monocytic origin. LC precursors are responsive to TGFβ, and we show that IL-34 specifically induces Id2 expression and survival of BM-derived cells.

### Immune damage and loss of eLC opens the epidermal compartment

Having considered LC repopulation at the cellular level, and demonstrated that monocytes differentiate into *bona fide* LC within the epidermis, we now considered the impact of immune pathology on the LC network and integrity of the epidermal compartment.

The kinetics of LC repopulation demonstrated that monocytes failed to completely replenish the LC network in most mice 10 weeks after transplant, suggesting a prolonged reduction in LC density. However, this decrease in LC numbers was also evident in mice that had received BMT alone, demonstrating that the slow-rate of division by mature LC ultimately dictated the speed at which the network was repaired, rather than LC origin. (Figure 7A). Confocal analysis of epidermal sheets revealed significant heterogeneity in the density of mLC in different fields of view (Figure 7B), but mLC tended to be smaller than eLC from BMT controls (Figures 7C). Notably, while mLC and dendritic epidermal T cells (DETC) were closely co-located within the epidermis, we found no difference in the frequency of mLC 3 weeks after BMT into *Tcrbd*^−/−^ recipients that lacked all endogenous T cells (Figure S9).

**Fig. 7.**
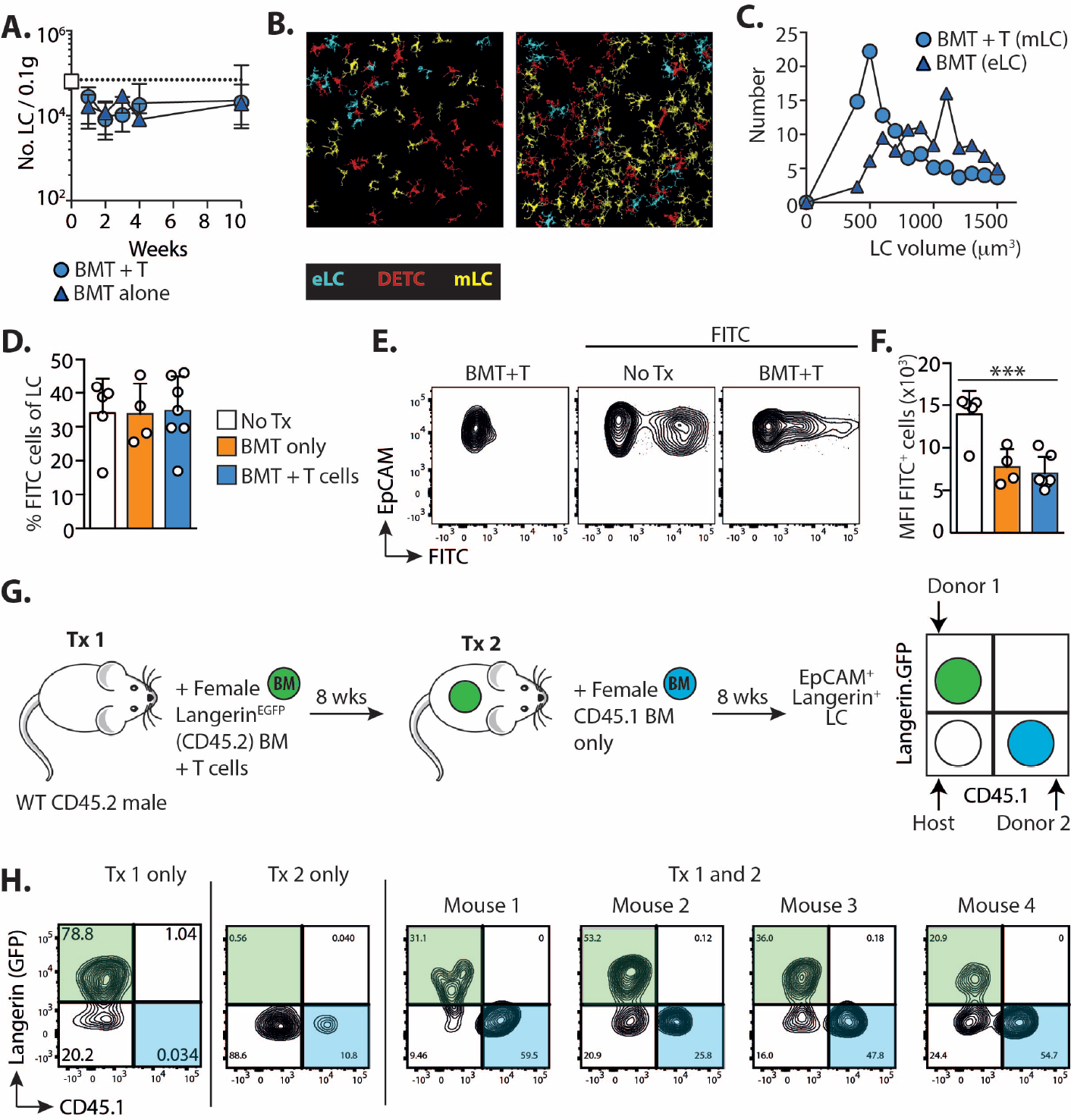
Immune damage and loss of eLC opens the epidermal compartment. Male mice received BMT with or without T cells. **A**. Graph shows the number ± SD of total CD11b^+^Langerin^+^ LC in mice receiving BMT alone (triangles) or BMT with T cells (circles). Data are pooled from 2 independent experiments (n=5-13). The white square and dotted line shows LC numbers ± SD in untreated controls (n=5). **B-C**. Epidermal sheets were stained with anti-Langerin and anti-CD45.2, and confocal images processed and quantified using the Definiens Developer software: eLC (cyan) are Langerin^+^CD45.2^+^; DETC (red) Langerin^neg^CD45.2^+^; and mLC (yellow) are Langerin^+^CD45.2^neg^. B. shows example images and the graph in C. is the volume of mLC compared to eLC from BMT controls. Data are from 1 transplant experiment with 3 BMT (20 fields of view analysed) and 2 BMT+T cell recipients (14 fields of view analysed) (n= 162 cells from BMT mice and 356 LC from BMT+ T cell recipients). **D.-F**. Topical FITC was painted on the ear skin of control un-transplanted mice (No Tx), or BMT and BMT + T cell recipients 10 weeks post-transplant. 3 days later uptake of FITC was analysed within MHCII^high^EpCAM^+^Langerin^+^ LC in draining LN. D. Bar graph showing the frequency ± SD of FITC^+^ cells within LC. E. Representative contour plots show FITC uptake in gated LC. F. Bar graph showing the FITC median fluorescent intensity ± SD within FITC^+^ LC. Data are pooled from 2 independent experiments (n=4-7), significance was analysed using a 1-way ANOVA, P=.0004. **G**. BMT + T cell recipients received a second round of irradiation and BMT alone 8 weeks later. The schematic illustrate the experimental setup. **H**. Flow plots show the outcome in the epidermis of independent mice, who have received the 1st transplant only (Tx 1), the 2nd transplant only (Tx 2) or both transplants (Tx 1 and 2). Contour plots are pre-gated on EpCAM^+^Langerin(PE-labelled)^+^ LC.

Given the smaller volume of mLC, we considered whether they were less integrated within the epidermis than eLC, and therefore migrated more readily in draining LN. To test this we topically applied FITC to the ear of mice which had received BMT 10 weeks earlier, with or without T cells. LN cells were divided into migratory and resident populations based on expression of CD11c and MHC II as published by others [40, 54, 55] and we determined the number of EpCAM^+^Langerin^+^ LC in the migratory gate (Figure S10). The frequency of FITC^+^ cells within LN LC was equivalent irrespective of LC origin (Figure 7D), demonstrating that, while mLC acquired the capacity to migrate to LN, this was not to a greater extent than eLC. It was evident from flow cytometry plots that mLC picked up less FITC that eLC from un-transplanted controls. Comparison of the median intensity of FITC^+^ cells demonstrated that this was indeed indeed the case, but a similar effect was also observed by eLC in BMT controls (Figure 7 E and F). Thus, mLC migrate to LN, but irradiation and BMT may also impact on the acquisition of topical antigen by LC. We next considered how changes to the density of the LC network would impact on the entry of cells into the epidermis in absence of T cell-mediated injury. Thus, transplanted mice received a second round of total body irradiation with BMT (without T cells) and we tracked the origin of epidermal LC 8 weeks later (Figure 7G). eLC were replaced by mLC after BMT with T cells (Tx1 only), but not BMT alone (Tx2 only), as expected. However, the epidermis of mice that had received both transplants (Tx1 and 2) contained 3 populations of co-existing LC. These were identified as radio-resistant eLC and mLC (Figure 7H Langerin.GFP^neg^CD45.1^neg^ and Langerin.GFP^+^CD45.1^neg^ respectively) and new BM-derived LC (Langerin.GFP^neg^CD45.1^+^).

In summary, mLC become long-lived quiescent LC that acquire the ability to self-renew, and migrate to draining LN. However, immune injury and destruction of eLC unseals the epidermal compartment and leads to long-term influx of new cells into the LC network.

## Discussion

We show that a single wave of monocytes is sufficient to replenish the LC network after T cell-mediated killing of eLC. Thus immune-pathology in the skin leads to functional adaptation of cellular mechanisms to ensure repopulation of the LC niche akin to those used to maintain resident macrophage populations in other tissues.

mLC become quiescent, self-renewing cells that acquire the capacity to migrate to draining LN. Moreover, monocytes differentiate into mLC are transcriptionally homologous to the eLC they replace, despite on-going T cell-mediated immune pathology in the epidermis. This finding was unexpected since monocytes that differentiate within other inflamed tissues remain transcriptionally distinct from their resident macrophage counterparts (33, 34), and microglia, which closely resemble LC in terms of their capacity to self-renew without contribution from circulating monocytes, are replaced by cells that retain a persistent monocytic signature (32).

Tissue-resident macrophages, including LC, are seeded from embryonic precursors before birth (5). However, while there is a clear consensus on the role of adult monocytes in maintaining and replenishing tissue macrophages in other organs, the nature of the precursor that repopulates LC in the skin has remained elusive. Previously, studies that addressed the nature of LC replacement in the epidermis have depended on the destruction of resident eLC by acute (15-30 minutes) exposure of ear skin to UV irradiation. However, these studies have produced conflicting data on whether Gr-1^+^ monocytes persist in the epidermis of UV-treated mice to become LC (17, 18). This work has led to the concept that an alternative ‘long-term’ LC precursor was required to replenish the LC network after UV-induced damage, and studies using human cells have since invoked a role for blood DC as LC precursors (24, 25). By contrast, we have used a model of allogeneic HSCT, in which donor T cells kill host eLC, and BM cells are recruited to the epidermis over a period of 4 weeks. Under these conditions monocytes can become longterm LC, and DC precursors do not contribute to the emerging LC network. It is conceivable that transient exposure to UV irradiation compared to the prolonged inflammation caused by allo-reactive T cells may trigger different mechanisms of LC repopulation in the skin. However, the adoptive transfer experiments previously used to define monocytes as LC precursors after UV irradiation are challenging, and require injection of large numbers of cells into *Ccr2*^−/−^*Ccr6*^−/−^ mice to reduce competition from endogenous cells (17, 18). We suggest that the physiological recruitment of monocytes from the BM in our model has revealed their role in the repair of the damaged eLC network.

We have demonstrated the dissociation of EpCAM and Langerin expression by LC precursors, markers previously thought to be expressed concomitantly only by mature cells (43). EpCAM^+^ cells are highly transient, but expressed a unique gene profile that clearly identifies them as distinct from both blood monocytes and mLC. Statistical analysis of our mathematical models clearly favours the linear differentiation of CD11b^high^ monocytes into EpCAM^+^ precursors of mLC. Nonetheless, this conclusion is specific to the models that we considered; the transient nature of the EpCAM^+^ population together with the similar kinetics of CD11b^high^ and EpCAM^+^ cells mean that it remains possible that the branched pathway may also occur with EpCAM^+^ cells as a developmental endpoint. However, we think this is unlikely; EpCAM^+^ cells express intermediate levels of CD11b, CD24 and DEC205, and the LDTF Id2 and Runx3 compared to CD11b^high^ monocytes and mLC, and EpCAM^+^ cells up-regulate Langerin upon culture *ex vivo*.

EpCAM^+^ cells express a specific panel of MMP, including *mmp14* (49). Spatial restriction of integrins avβ6 and avβ8 to hair follicle keratinocytes controls residency of eLC by activation of latent TGFβ (56). Moreover, human monocytes have recently been shown to employ the combination of MMP14 and αvβ8 to activate latent TFGβ in the gut (57). We postulate that homotypic binding to EpCAM^+^ keratinocytes, and secretion of MMP, permits residency and local conditioning of the epidermal niche by EpCAM^+^ cells, before differentiation into mLC. However, despite the surge of monocytes entering the epidermis, and proliferation of CD11b^high^ cells *in situ*, we have identified bottleneck with only 4% of CD11b^high^/EpCAM^+^ cells up-regulating Langerin to become mLC. It is not clear whether similar inefficiencies also occur for monocyte-derived macrophages at other sites. Notably, monocyte-derived EpCAM^+^Langerin^neg^ cells can be identified within the oral mucosa, wherein inflammation blocks the transition to mucosal LC (13, 58). Likewise, activated monocytes (59, 60) accumulate within psoriatic epidermis and contribute to disease pathology. Therefore, it is possible that continued inflammation in our model blocks differentiation of EpCAM^+^ cells, which may also directly contribute to immune-pathology in diseased skin.

Cell cycle genes are switched off in circulating monocytes, but differentiation into mature tissue-resident macrophages is associated with the activation of a self-renewal program (61). Entry of CD11b^high^ cells into the epidermis triggers a burst of proliferation that has also been reported when phagocytic monocytes enter the skin after UV-irradiation (17). Subsequent to this, mLC continue to divide at homeostatic levels. Strikingly, our mathematical modelling data predict a turn-over rate for mLC 10 weeks post-transplant that is remarkably similar to published estimates for steady state eLC (12, 51). This estimate not only strongly supports the accuracy of our fits, but further reinforces the developmental convergence of quiescent mLC with their embryonic counterparts. It has been suggested that the density of tissue macrophage populations is controlled by mechanisms of quorum sensing in response to CSF-1 (62), but this has not been directly demonstrated experimentally. We found that models which assumed constant rates of proliferation and loss were not sufficient to explain the waning of the number of Ki67^+^ LC over time, and density-dependent proliferation was required to fit the data. Thus, our findings strongly support the concept of quorum sensing within the epidermal niche. CSF1 and IL-34 compete for the CSF1 receptor (63). Given the dominance of IL-34, rather than CSF1 in the epidermal environment (64), and our *in vitro* data showing that IL-34 increases expression of *Id2* and promotes BM-LC survival, it is possible that IL-34 fulfils this function in the skin.

The mechanisms by which immune cells enter the epidermis have not been fully elucidated, but monocytes are excluded in the absence of injury, possibly due to competition with resident cells (62). We have shown that the epidermal compartment is not resealed after immune-mediated destruction of eLC, and BM-derived cells continue to be recruited into the epidermis. It is notable that small numbers of donor CD11b^+^ cells constitutively enter the epidermis of transplanted mice in the steady state, as observed 10 weeks post-transplant (see Figure 3D). One possibility is that the inefficient repair of the mLC network reduces competition between established and incoming cells, however the signals that continue to recruit monocytes across the barrier membrane and into the steady state epidermis remain to be defined. Based on these data we suggest that immune pathology permanently opens the epidermal compartment, leading to long-term heterogeneity within the LC network as monocytes continue to be recruited into the skin. In this sense, the LC network in the epidermis more closely resembles that of the oral mucosa (1, 13) after exposure to immune injury.

The activation of auto- or allo-reactive T cells, and destruction of tissue-resident cells can have profound impacts on the balance of immune cells within tissue compartments with long-term consequences for the control of infection and cancer at these sites. The skin is highly sensitive to such changes and is a major target organ for T cells in patients suffering from graft-versus-host disease following HSCT, and those receiving immune checkpoint blockade (65). However, we know little about the impact of immune injury in these patients on the regulation of immunity in the skin. Here we provide novel insights into the cellular mechanisms by which the LC compartment is replenished and maintained after damage, and demonstrate that immune injury triggers an adaptive process that converges closely with mechanisms to regenerate other tissue-resident myeloid cells.

## Materials and methods

### Mice

C57BL/6 (B6) mice were purchased from Charles River and bred in house by the UCL Comparative Biology Unit. Langerin.DTR.EGFP mice (66) were originally provided by Adrian Kissenpfennig and Bernard Malissen. B6 TCR-transgenic anti-HY MataHari (Mh) mice (67) were provided by Jian Chai (Imperial College London, London, UK). Ccr2 knock out mice were a gift from Frederic Geissmann (68). All strains were bred in house at UCL. *TCRbd*^−/−^ mice were generated by crossing *Tcrd*^−/−^ (69) and *Tcrb*^−/−^ (70) lines, and bred in house at Imperial College London Hammersmith campus. CD45.1 OT-I TCR transgenic mice were bred in house. All procedures were conducted in accordance with the UK Home Office Animals (Scientific Procedure) Act of 1986, and were approved by the Ethics and Welfare Committee of the Comparative Biology Unit, Hampstead Campus, UCL, London, UK.

### Bone marrow transplants

Recipient male CD45.2 B6 mice were lethally irradiated (11 Gy total body irradiation, split into 2 fractions over a period of 48 hours) and reconstituted 4 hours after the second dose with 5×10^6^ female B6 CD45.1 T cell-depleted BM cells and 2×10^6^ CD4 T cells, with 1×10^6^ CD8 Thy1.1^+^ Mh T cells, administered by intravenous injection through the tail vein. T cells were isolated by magnetic selection of CD4 or CD8 BM cells or splenocytes using the Miltneyi MACS system (QuadroMACS Separator, LS columns, CD4 [L3T4] MicroBeads, CD8a [Ly-2] MicroBeads; Miltenyi Biotec) according to the manufacturer’s instructions, either to be discarded from T cell-depleted BM, or selected for injection of splenic T cells. In some experiments Langerin.DTR.EGFP recipients or BM donors were used to track Langerin^+^ host or donor LC respectively.

For secondary irradiation experiments, BMT recipients initially received BM from Langerin.DTR.EGFP donors with CD4 and CD8 Mh T cells. 8 weeks later, mice received 11 Gy split dose total body irradiation and female B6 CD45.1 T cell-depleted BM alone.

### Mixed chimera experiments

BM from Vav-Cre.Rosa26LSL^Tomato^ and Clec9a-Cre.Rosa26LSL^YFP^ donors was a gift from Caetano Reis e Sousa (Francis Crick Institute). Irradiated B6 male mice received a 50:50 mix of BM from the reporter mice with T cells. For CCR2 competitive chimeras, irradiated CD45.2^+^ Langerin-DTR.EGFP recipients received a 50:50 mix of female CD45.2 *Ccr2*^+/+^ and CD45.1 *Ccr2−/−* BM with T cells.

### FITC painting

The dorsal side of ear pinna were coated with 25µl of a 1:1 mixture of 0.5% FITC (Fluorescein isothiocyanate, Sigma-Aldrich, UK) in acetone and dibutyl phthalate (Sigma-Aldrich, UK). 72 hours later the draining auricular and cervical LN were harvested and analysed.

### Bone marrow cultures

BM was flushed from femurs and tibias of donors, and red blood cells lysed in 1ml ammonium chloride (ACK buffer, Lonza UK) for 1 minute at room temperature. Cells were washed and resuspended at 2.5×10^6^/ml in R5 medium (RPMI 1640, Lonza, Switzerland), 5% heat-inactivated FBS (Life Technologies, USA), 1% L-glutamine (2 mM; Life Technologies, USA), 1% Pen-strep (100 U/ml Life Technologies, USA) and 50µM β-ME (Sigma-Aldrich, UK). 1ml of cell suspension was plated per well in tissue culture-treated 24-well plates and supplemented with 20ng/ml recombinant GM-CSF and 5ng/ml TGFβ (Peprotech, USA). Cells were cultured at 37°C. Media was partially replaced on day 2 of culture, completely replaced on day 3, and the cells harvested on day 6. Some cultures were supplemented with combinations of 8µg/ml IL-34 (Generon, UK), 100 µg/ml BMP7 (RD systems, USA and 10µg/ml CSF-1 (Biolegend, USA) for the duration of the culture.

### Flow cytometry

Cells were distributed in 96 well conical bottom plates and incubated in 2.4G2 hybridoma supernatant (containing αCD16/32) for at least 10 min at 4°C to block Fc receptors. For cell surface labelling, cells were incubated with fluorochrome-conjugated antibodies diluted in 100µl FACS buffer (PBS/1mM EDTA/1% FBS) at 4°C for at least 20 min in the dark: EpCAM (G8.8, eBioscience, USA), CD11b (M1/70, eBioscience, USA), CD45.1 (A20, BD Biosciences, Germany), CD45.2 (104, eBioscience, USA), MHC II I-A/I-E (M5/114.15.2, eBioscience, USA), CD11c (HL3, BD Pharmingen, USA), CD24 (M1/69 BD Biosciences or Biolegend), CD205 (205yekta, eBioscience), CD115 (AFS98, eBioscience), CD135 (A2F10, eBioscience), B220 (RA3-6B2, BD Biosciences), Vβ8.3 TCR (1B3.3, BD Biosciences, Germany). To exclude lineage^+^ cells, we used a cocktail of CD3 (145-2C11, BD Biosciences, Germany), CD19 (1D3BD Biosciences, Germany), NK1.1 (PK136, Biolegend) and Ly6G (1A8, BD Biosciences, Germany) all conjugated to APC-Cy7.

Intracellular staining with αLangerin (CD207) antibodies (eBioL31, eBioscience, USA) was performed after cell surface immunolabelling. Samples were washed with FACS buffer, fixed in 100µl fixation solution (BD Cytofix/Cytoperm solution, BD Biosciences, UK) for 15 min at 4°C, washed twice with permeabilization buffer (BD Perm/Wash, BD Biosciences, UK) and incubated with 100µl αLangerin diluted in permeabilization solution at 4°C for 30 min in the dark.

Live cells were identified by exclusion of propidium iodide (unfixed cells) (Life Technologies, USA), or a fixable viability dye (eBioscience, USA or Life Technologies, USA). Multicolor flow cytometry data were acquired with BD LSR-Fortessa and BD LSR II cell analyzers equipped with BD FACSDiva v6.2 software (BDBiosciences, Germany). Fluorescence activated cell sorting was performed on a BD FACSAria equipped with BD FACSDiva v5.0.3 software (BD Biosciences, Germany). All samples were maintained at 4°C for the duration of the sort. Cells were sorted into PBS/1% FBS before resuspension in Buffer RLT (QIAGEN, USA) or directly into Buffer RLT with 1% 2-β-mercaptoethanol (Sigma, UK), disrupted through vortexing at 3200 rpm for 1min, and immediately stored at −80°C until further processing. 2-3 biological replicates were obtained for each sample from at least 2 independent experiments, each containing a minimum of 4000 cells (pooling where necessary from multiple mice from individual experiments). Flow cytometry data were analyzed with FlowJo X v9 and 10 (LLC, USA), and cells were pre-gated on singlets (FSC-A versus FSC-H), and a morphological FSC/SSC gate.

### Measurement of proliferation

*In vivo*. Epidermal single cell suspensions were immuno-labelled with surface antibodies, then fixed and permeabilized using the eBioscience intranuclear staining kit, before incubation with αKi67-v450 antibodies (SolA15, eBioscience, USA). Gates were set on non-proliferating cells and unstained cells. Alternatively, mice were injected with 100µg 5-Ethynyl-2′-deoxyuridine (EdU) i.p (Invitrogen, USA) and euthanized 4 hours later. *In vitro*. Cells were pulsed with 10µM EdU on day 2 or day 5 of culture and the medium replaced 24 hours later. For both *In vivo* and *In vitro* studies, cells were labelled for flow cytometry using the Click-iT Plus EdU Flow Cytometry Assay Kit (Invitrogen, USA), according to the manufacturer’s instructions.

### Immunohistochemistry

4mm biopsy punches were excized from the dorsal and ventral sides of split ears and incubated in 0.5M ammonium thiocyanate for 30 min at 37°C to remove the epidermis. Epidermal sheets were collected in eppendorf tubes, washed twice with PBS, and fixed with cold (stored at −20°C) acetone for 10 min. Sheets were washed twice with PBS, and blocked using 0.25% fish gelatin, 10% normal goat serum in PBS for 1 hour at room temperature. Sheets were then incubated with primary rat αLangerin (eBioscience), and αCD45.2-biotin (eBioscience) antibodies (both diluted 1:100 in blocking buffer), and incubated for 1 hour at room temperature. Sheets were washed 4 times in PBS, and incubated with goat αRat-Alexa 647 (Jackson ImmunoResearch Laboratories, 1:1000) and Streptavidin-e570 (eBioscience, 1:200) secondary antibodies diluted in blocking buffer for 1 hour at room temperature. Stained sheets were washed 4 times with PBS, and mounted on slides with ProLong™ Diamond anti-fade mountant (Invitrogen). Samples were imaged on a Nikon Ti inverted microscope, through a 20X objective (Plan Apochromat N.A. 0.75 W.D. 1mm) or 40X objective (Plan Apochromat N.A. 0.95 W.D. 0.21mm), using a C2 confocal scan head with488nm and 561/568nm optimized fluorescence filter cubes (Nikon Instruments, Tokyo, Japan). Multiple Z-stacks were acquired for each sample. Data was saved as nd2 files using FIJI/ImageJ for quantification.

### Confocal analysis

Quantification of confocal records was performed using Definiens Developer software. Each channel in a record was processed with Gaussian filter followed by application of multi-resolution segmentation. Individual cells were detected based on their relative intensity in Langerin and CD45.2 channels, and assigned a colour according to their identification as Langerin^+^CD45.2^+^ eLC, Langerin^+^CD45.2^neg^ dLC or Langerin^neg^CD45.2^+^ DETC. The cell volume (µm^3^ based on total number of voxels occupied by a cell).

### Statistics

All data, apart from RNAseq data, were analysed using GraphPad Prism Version 6.00 for Mac OsX (GraphPad Software, USA).

## ACKNOWLEDGEMENTS

We thank the UCL Comparative Biology Unity for their support with animal work, and UCL Genomics for performing the RNA sequencing. We are grateful to Caetano Reis e Sousa for providing the Clec9a reporter mice, to Joe Grove and Benedict Seddon for constructive discussions during this work, and Hans Stauss for his support. This work was funded by a BBSRC project grant (BB/L001608/1) and Royal Free charity funding to C.L.B., an MRC PhD studentship (1450227) to H.W. and the NIH (R01 AI093870) for A.J.Y. D.S.U. acknowledges the support of the Wellcome Trust (grant 100156), and the Infection and Immunity Immunophenotyping (3i) Consortium.

The authors declare no competing interests.

Conceptualization, C.L.B.; Methodology, I.F., H.W., A.J.Y. and C.L.B.; Investigation, I.F., H.W. and P.S.; Formal analysis, S.H. D.S.U. and A.J.Y.; Resources, J.S.; Writing – Original Draft, C.L.B and A.J.Y; Writing – Review and Editing, I.F., H.W, J.S., R.C, and C.L.B; Funding Acquisition and supervision, C.L.B.

