## Supplemental material for "A single wave of monocytes is sufficient to replenish the long-term Langerhans cell network after immune injury"

**Supplemental materials**

Figure S1. Schematic showing the BMT protocol.

Figure S2. Gating of BM-derived cells.

Figure S3. Sorting strategies used to isolate cells for RNAseq.

Figure S4. EpCAM$^+$ cells are intermediates between blood monocytes and donor LC.

Figure S5. Modelling the cellular dynamics of epidermal myeloid cells.

Figure S6. Enrichment of innate signalling and T cell activation pathways in dermal monocytes.

Figure S7. Host eLC show responses to the inflammatory environment post-BMT.

Figure S8. mLC rapidly differentiate in the epidermis, but host eLC show responses to the inflammatory environment post-BMT.

Figure S9. Repopulation of DETC-deficient hosts by LC.

Figure S10. Gating strategy to identify migrating LC.

Table S1. Differentially expressed genes between eLC and mLC.

Supplemental methods including RNA sequencing and mathematical modelling.

Supplemental references.

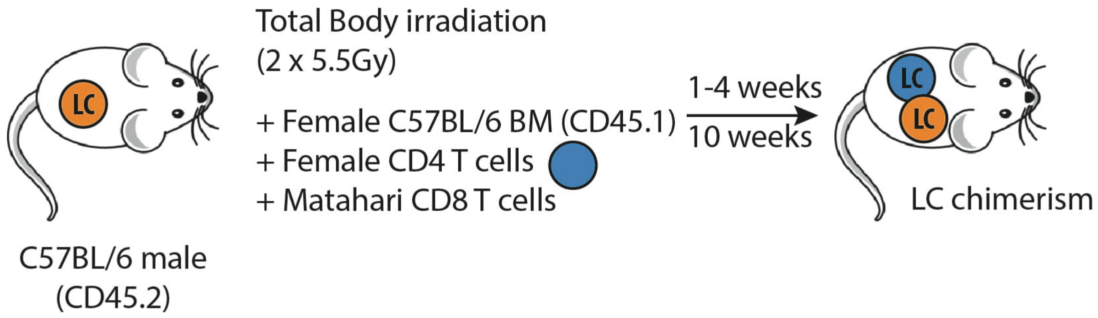


**Figure S1. Schematic showing the BMT protocol.** Male CD45.2 C57BL/6 mice were irradiated and received congenic (CD45.1) BM and polyclonal CD4 T cells, with $10^6$ CD8 Matahari T cells. Recipients were analysed at different time points for LC chimerism.

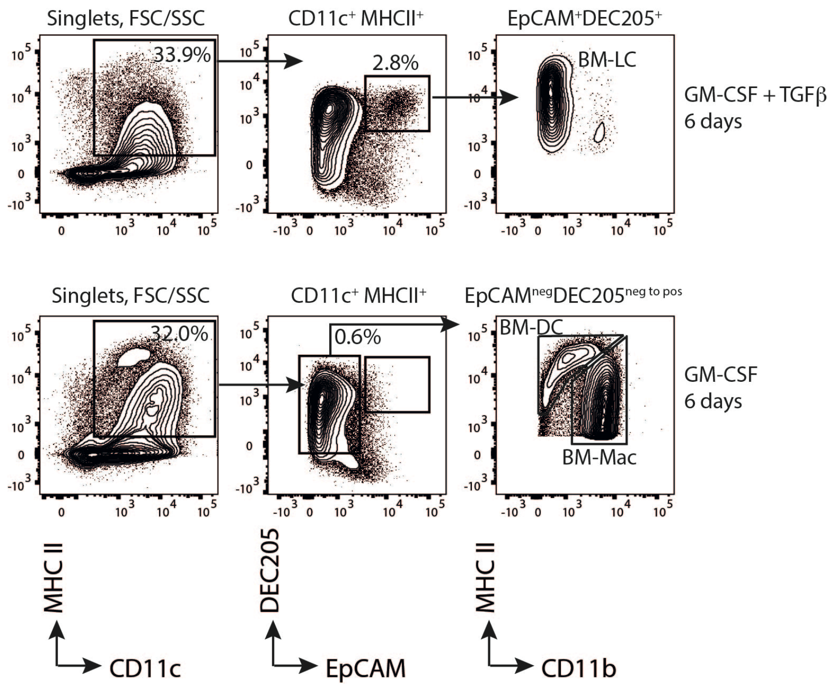


**Figure S2. Gating of BM-derived cells.** Murine C57BL/6 BM was cultured for 6 days with GM-CSF alone, or with TGFβ. Representative contour plots show the phenotype of cells on day 6 of culture. EpCAM$^{+}$DEC205$^{+}$ BM-LC differentiated in cultures that contained TGFβ. Alternatively, GM-CSF alone produced DEC205$^{neg\ to\ pos}$ cells that were sub-divided into MHCII$^{high}$CD11b$^{low}$ BM-DC and MHCII$^{int\ to\ low}$CD11b$^{high}$ BM-macrophages (mac).

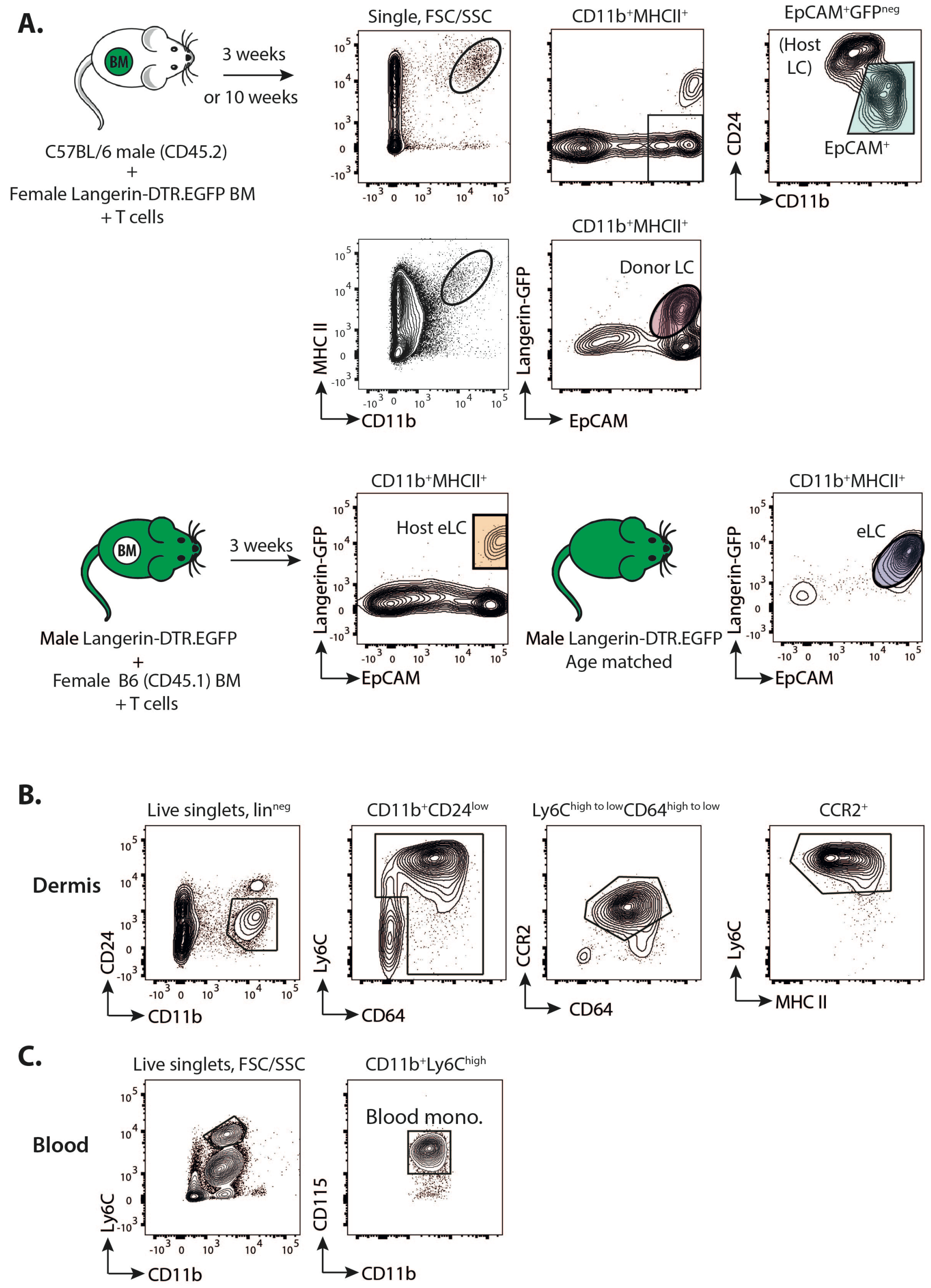


**Figure S3. Sorting strategies used to isolate cells for RNAseq. A.** Schematic showing the transplant strategy using Langerin.DTR.EGFP BM to identify and sort EpCAM$^+$ cells and LC. Flow plots show pre-gated single epidermal cells. *Top* - donor EpCAM$^+$ cells were identified by gating on CD11b$^{high}$CD24$^{low}$ cells within the CD11b$^+$MHCII$^+$EpCAM$^+$Langerin.GFP$^{neg}$ gate. Alternatively, donor LC were identified either 3 weeks or 10 weeks post-transplant by expression of EpCAM and Langerin.GFP. *Bottom* - Male Langerin.DTR.EGFP mice received female congenic bone marrow. Host eLC were isolated 3 weeks later based on co-expression of EpCAM and Langerin.GFP. Alternatively, untransplanted, age-matched male Langerin.DTR.EGFP mice were use to sort control LC populations. **B.** Contour plots show the gating strategy used to sort activated monocytes from dermal single cell suspensions. Cells were identified as CD11b$^+$CD24$^{neg}$CCR2$^+$Ly6C$^{high}$MHCII$^{low\ to\ +}$, as described by Tamoutounour *et al.* **C.** Contour plots showing gating of CD11b$^+$Ly6C$^{high}$CD115$^+$ blood monocytes.

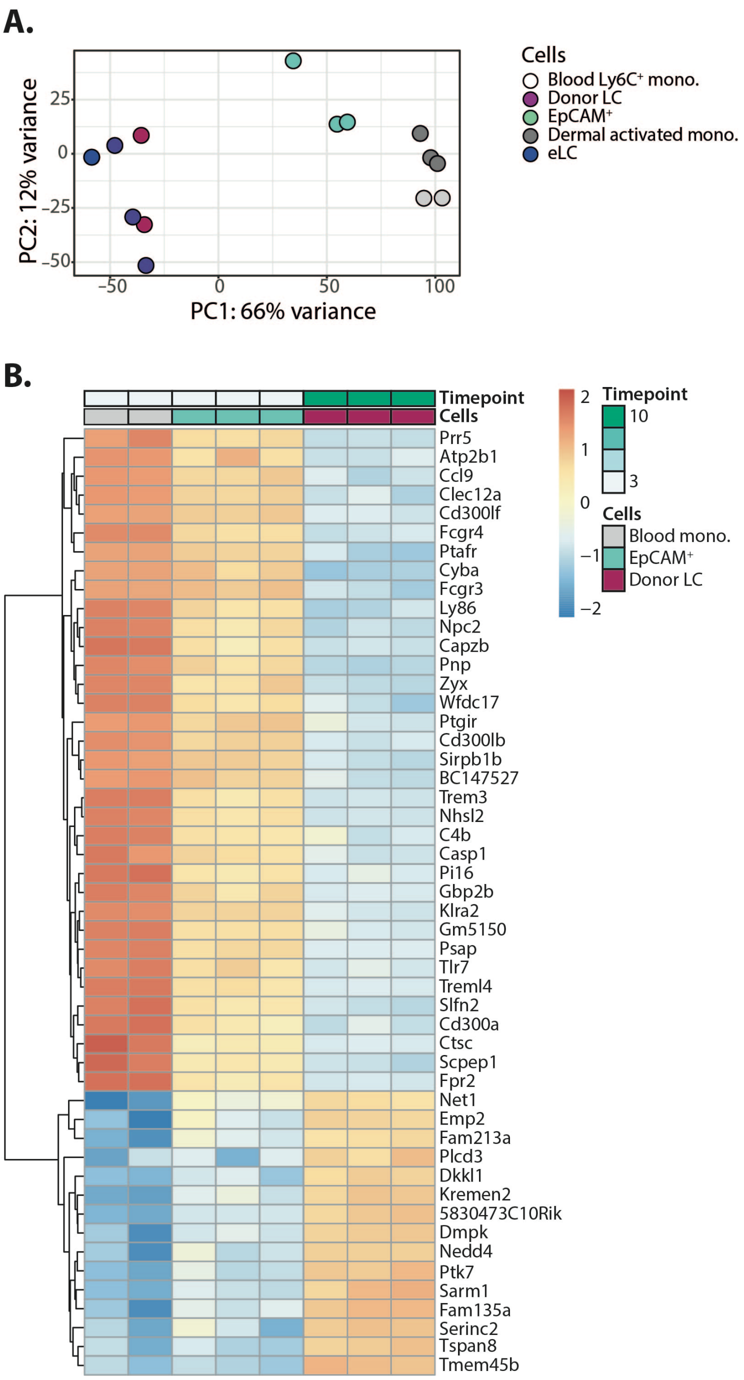


**Figure S4. EpCAM⁺ cells are intermediates between blood monocytes and donor LC. A.** Principal components analysis of the different cell populations. 66% of the variance within the expression data is orthogonal to PC1. **B.** Heat map showing the relative expression of the top 50 genes that contributed to the differences between blood monocytes, EpCAM⁺ cells (3 weeks post-transplant) and donor LC (10 weeks post-transplant) along the PC1 axis.

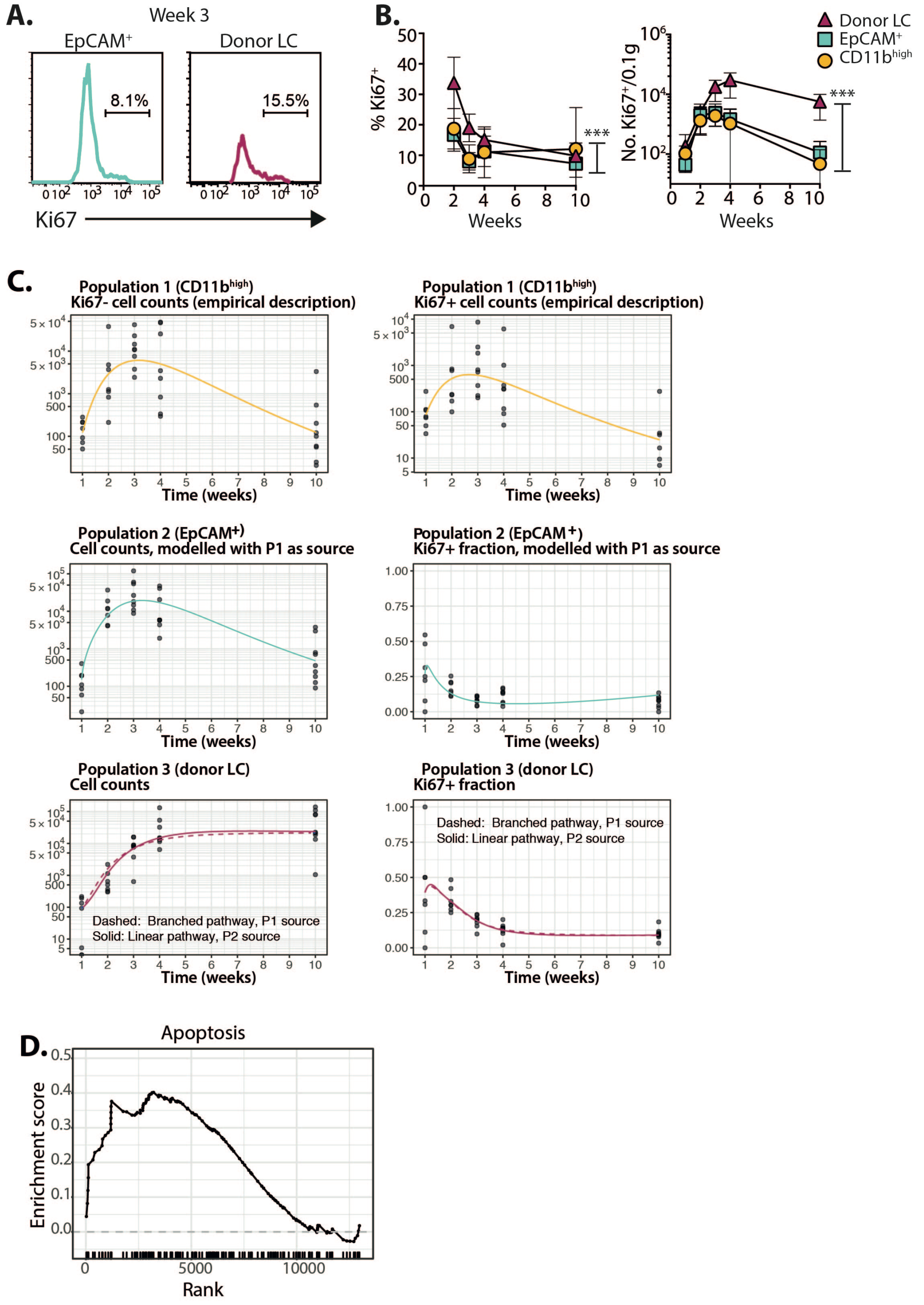


**Figure S5. Modelling the cellular dynamics of epidermal myeloid cells. A.** Representative histograms show gating of Ki67$^{+}$ cells in the EpCAM$^{+}$ and donor LC populations 3 weeks after BMT with T cells. **B.** Graphs show the frequency ± SD (*left*) and number ± SD per 0.1g total ear tissue (*right*) of Ki67$^{+}$ cells within gated epidermal populations. Circles CD11b$^{high}$; squares EpCAM$^{+}$; triangles donor LC. P<.0001, with 2-way ANOVA. Data are pooled from 2 independent experiments per time point (n=7-8). **C.** Data from the experiments shown in B. were modelled mathematically. *Upper panels* - Fitted, empirical descriptions of the timecourses of Ki67$^{+}$ and Ki67$^{neg}$ CD11b$^{high}$ cells (P1). *Middle panels* - fits to the total numbers and Ki67$^{+}$ fraction of EpCAM$^{+}$ cells (P2), using the empirical descriptions of the CD11b$^{high}$ cell kinetics as a source. *Lower panels* - fits to timecourses of mature donor LC (P3) numbers and Ki67$^{+}$ fraction using either CD11b$^{high}$ (P1) or EPCAM$^{+}$ cells (P2) as a source. Details of models are given in Supplemental Materials and Methods. **D.** Plot from a gene set enrichment analysis comparing EpCAM$^{+}$ cells to donor LC at week 10, showing enrichment of genes related to apoptotic pathways in EpCAM$^{+}$ cells. Normalized enrichment score 1.457, FDR q-value 0.024.

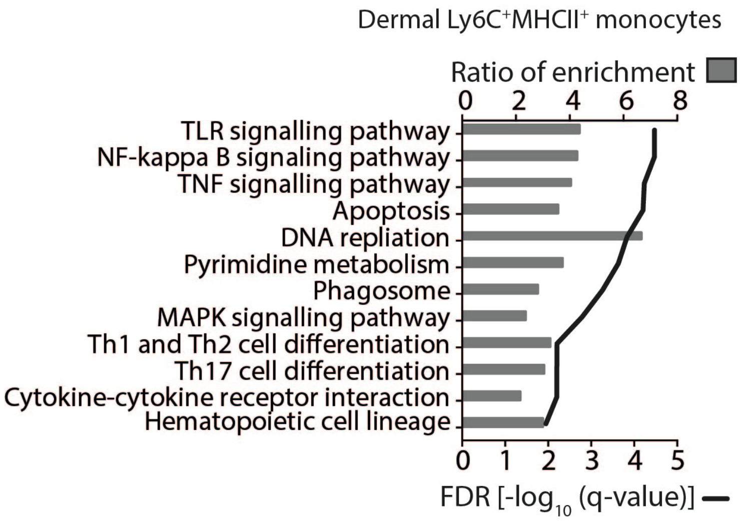


**Figure S6. Enrichment of innate signalling and T cell activation pathways in dermal monocytes.** The graph shows the ratio of enrichment (bars) and FDR *q* values (line) for pathways predicted by WebGestalt to be over-expressed by dermal Ly6C+MHCII+ activated monocytes compared to blood monocytes. Over-expressed genes had a fold change ≥ 2, $q < 0.05$.

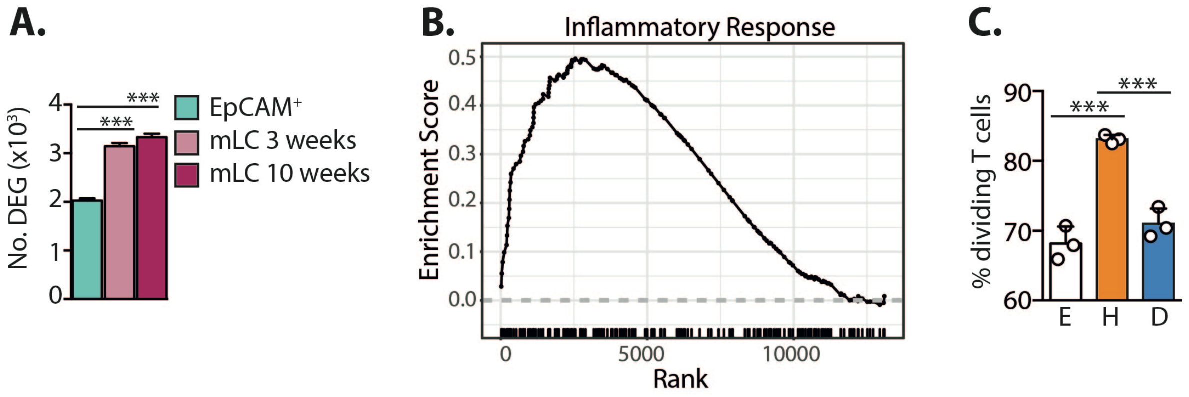


**Figure S7. mLC rapidly differentiate in the epidermis, but host eLC show responses to the inflammatory environment post-BMT.** **A.** Bar chart showing the number of differentially expressed genes (DEG) in each population compared to blood Ly6C$^+$ monocytes. EpCAM$^+$ vs donor LC at 3 or 10 weeks, one-way ANOVA with Holm-Sidak's test $P$<.0001. **B.** Plot from gene set enrichment analysis comparing host eLC to donor LC at week 3, showing enrichment of genes related to the inflammatory response in host eLC. Normalized enrichment score 1.438, FDR q-value 0.064. **C.** Host (H) or donor (D) LC were sorted from the epidermis 3 weeks post-BMT + T cells, or eLC isolated from untreated controls. Langerin.DTR.EGFP mice were used to identify host or donor cells, as described in Figure S3, and co-cultured with antigen-specific CD8 T cells. The graph compares proliferation ± SD of for cultures with 100 LC per well. E versus H $P$=.0002, H versus D $P$=.0005, one-way ANOVA with Holm-Sidak's multiple comparisons test. Symbols are technical replicates, and data are representative of LC sorted in 2 independent experiments.

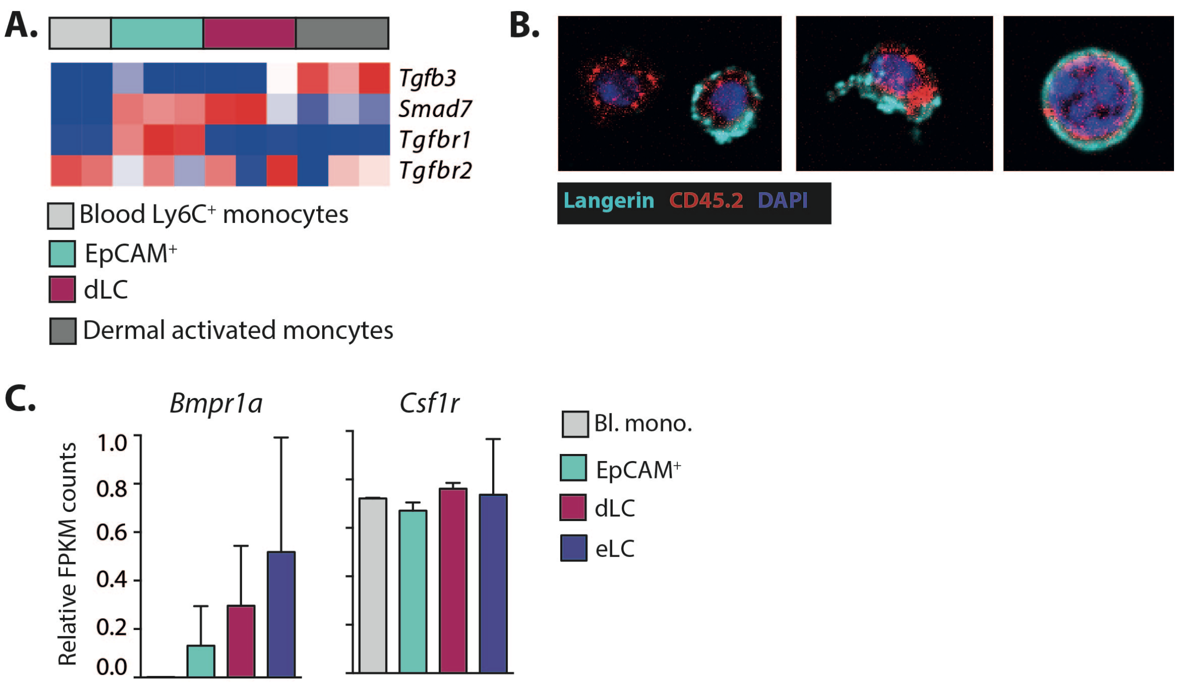


**Figure S8. EpCAM⁺ cells are responsive to epidermal growth factors. A.** Heat map showing the relative expression of genes associated with TGFβ signalling by different myeloid cell populations. **B.** Microscopy images of representative sorted EpCAM⁺ cells after culture with primary syngeneic keratinocytes and TGFβ for 36 hours. Cells were fixed and stained for CD45.2 and Langerin. Images are representative of cells from 2 independent experiments. **C.** Graphs show the relative gene expression (FKPM) of different receptors from the RNAseq data. Blood Ly6C⁺ monocytes n = 2, epidermal EpCAM⁺ cells n= 3, donor LC n = 3, age-age-matched host LC n = 3. There were no statistically significant differences between groups.

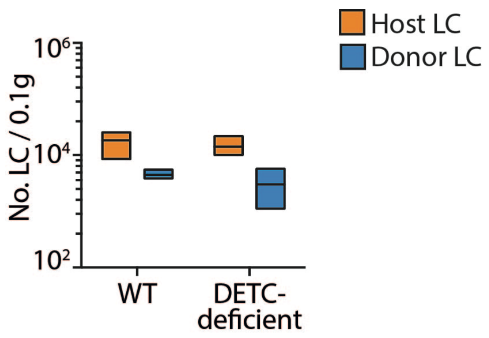


**Figure S9. Repopulation of DETC-deficient hosts by LC.** Bars show the number (mean and range) of host (orange) eLC or donor (blue) mLC in the epidermis of WT or DETC-deficient recipients 3 weeks after BMT with T cells. Data are from 1 representative experiment of 2 (n= 3-4).

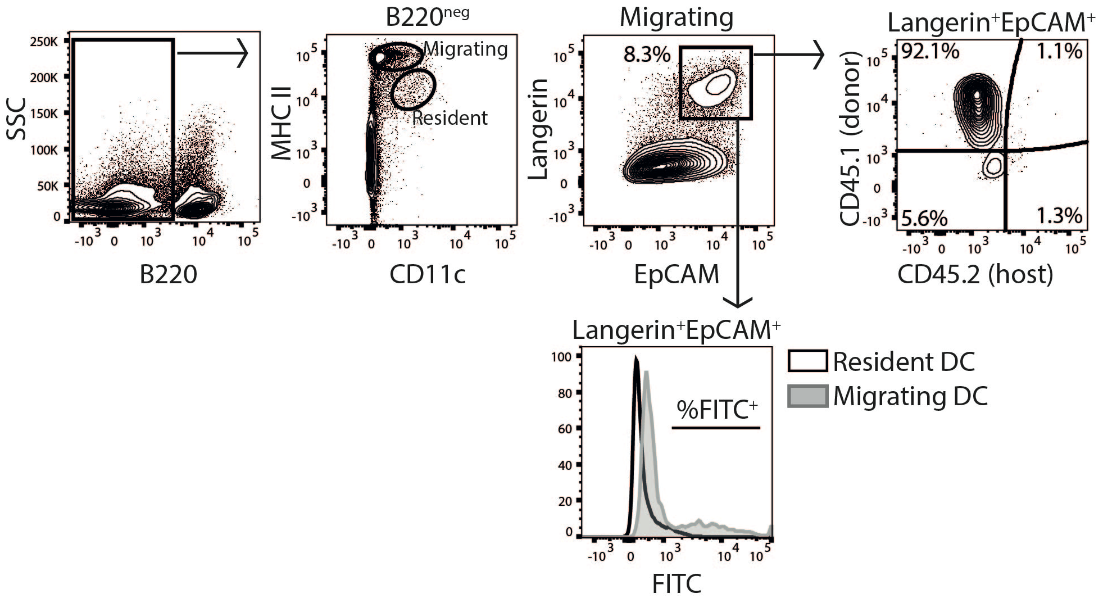


**Figure S10. Gating strategy to identify migrating LC.** FITC was painted onto the ears of control mice or those that had received BMT with or without T cells 10 weeks earlier. Representative plots show the gating strategy. B220$^{+}$ cells were excluded from live, single cells. Of the B220$^{neg}$ cells, populations were identified as CD11c$^{+}$MHCII$^{high}$ migrating cells and CD11c$^{high}$MHCII$^{+}$ resident cells. LC were identified within the migrating cells by co-expression of Langerin and EpCAM, and these cells were confirmed to be derived from donor CD45.1$^{+}$ BM. The histogram overlay shows that up-take of FITC is restricted to gated migrating cells.

| Base mean | Log2 fold change | P value | Adjusted P value | Gene name | Suggested fuction |
|---|---|---|---|---|---|
| 49.78040956 | -44.65292318 | 1.85E-47 | 5.41E-42 | ***Vcam1*** | **Cell adhesion** |
| 9.000189682 | -42.68969063 | 5.71E-25 | 8.36E-20 | ***Gnb3*** | G protein - signalling |
| 22.65879277 | -40.69285895 | 3.84E-23 | 9.54E-19 | ***Muc15*** | **Cell adhesion** |
| 279.4903999 | 3.170389599 | 6.59E-18 | 3.64E-14 | *Rps4l* | Ribosomal |
| 5429.312963 | -1.482841516 | 5.84E-09 | 3.41E-06 | ***Tmsb10*** | **Cytoskeleton organisation** |
| 499.9550931 | -2.48297942 | 4.83E-09 | 8.06E-06 | ***Acot7*** | Regulation of lipid metabolism |
| 739.1650105 | 1.708605491 | 3.76E-08 | 4.88E-05 | *Polg* | DNA polymerase subunit gamma |
| 3869.97247 | -1.386401031 | 2.04E-07 | 7.91E-05 | ***Stat1*** | Interferon signaling |
| 11.87569702 | 17.55448961 | 4.85E-08 | 1.10E-04 | *Heg1* | Mucin related? |
| 26268.18931 | -1.890512679 | 3.55E-07 | 1.10E-04 | ***Mfge8*** | **Cell adhesion** |
| 194.8187034 | 2.993004417 | 1.02E-08 | 1.58E-04 | *Rbfox1* | RNA-binding protein |
| 910.2978017 | -1.461432195 | 5.84E-07 | 4.42E-04 | ***Arhgap31*** | GTPase-activating protein |
| 12.18004708 | 17.80094552 | 2.88E-07 | 4.71E-04 | *Nme5* | Anti-apoptotic |
| 345.2468399 | -2.774360982 | 2.80E-07 | 4.71E-04 | ***Map4k1*** | JNK activation |
| 2358.320843 | -4.855243937 | 2.88E-06 | 5.19E-04 | ***Iigp1*** | Target for Stat1 signalling |
| 4533.48776 | 1.356125189 | 2.82E-06 | 5.19E-04 | *Ifitm2* | Interferon-induced gene |
| 913.9626163 | 0.663761668 | 2.07E-06 | 0.001175993 | *Unc50* | RNA-binding |
| 474.1602636 | -3.896334558 | 1.87E-06 | 0.001319679 | ***Cyfip2*** | **Cell adhesion** |
| 2524.360564 | -1.287896324 | 1.07E-05 | 0.001333934 | ***Clec4n*** | Dectin-2. Binds mannose-containing structures |
| 4191.506502 | -2.192436644 | 1.37E-05 | 0.001619773 | ***Fgl2*** | **Fibrinogen-like** |
| 540.1680501 | -0.880769708 | 3.65E-06 | 0.00242965 | ***Whsc1l1*** | Chromatin organisation |
| 1442.820134 | -1.276857164 | 2.34E-05 | 0.002811012 | ***Fmnl1*** | **Cell motility and survival in macrophages** |
| 3054.345797 | -0.929166308 | 2.63E-05 | 0.002811012 | ***Csf2rb*** | CSF-2 receptor β common sub-unit |
| 476.9476862 | -1.270764599 | 6.46E-06 | 0.002811012 | ***Cbfa2t3*** | Transcriptional corepressor with CBFA2/Runx1 |
| 1472.138632 | -1.440378933 | 3.15E-05 | 0.002990642 | ***Elovl5*** | Fatty acid synthesis |
| 545.4110615 | -1.411130891 | 1.06E-05 | 0.004073344 | ***Cep83*** | Organelle biogenesis |
| 421.2704847 | -3.897887201 | 1.65E-05 | 0.006079613 | ***Ifit1*** | Interferon-induced gene |
| 1057.869752 | -3.065446226 | 1.64E-05 | 0.006079613 | ***Oasl2*** | Type I IFN signalling |
| 370.5617155 | -3.578504542 | 7.98E-06 | 0.006079613 | ***P2ry6*** | G-protein-coupled receptor |
| 1161.58779 | -0.893424545 | 9.19E-05 | 0.007037555 | ***Gls*** | Glutaminase - metabolism |
| 522.6837969 | 0.686416805 | 1.72E-05 | 0.007037555 | *Dpm2* | Protein metabolism |
| 4.83174206 | 30.55808943 | 1.21E-07 | 0.007111331 | *Kcnma1* | Potassium channels, actin binding. |
| 1124.507266 | -1.276769384 | 8.14E-05 | 0.007111331 | ***Csf2rb2*** | Csf2rb pseudogene |
| 2177.362624 | -2.416907594 | 1.04E-04 | 0.00751129 | ***Lgals3bp*** | Galectin-binding |

**Table S1.** List of all differentially expressed genes between mLC 10 weeks post-BMT with T cells, and eLC from age-matched controls. Genes up-regulated in donor LC are highlighted in bold. Suggested functions are inferred from gene ontology terms on the GeneCard® database (https://www.genecards.org).

**Supplemental Materials and Methods.**

**Sample preparation for gene expression analysis.** RNA was extracted using the RNeasy Micro Kit (QIAGEN, USA) following the manufacturer's protocol.

*RT-PCR.* cDNA was synthesized using a QuantiTect Reverse Transcription Kit (Qiagen, USA). The quantitative reverse transcription PCR (qRT-PCR) was run on a CFX96 Touch Real Time PCR detection system (Bio-Rad, USA), using a Quantifast SYBR Green PCR kit (Qiagen, USA). Primer sequences for *Id2 (*F *AAAACAGCCTGTCGGACCAC* and R *CTGGGCACCAGTTCCTTGAG*) and *Runx3* (F *CAGGTTCAACGACCTTCGATT* and R *GTGGTAGGTAGCCACTTGGG*) were obtained from published sources (2, 3). mRNA expression was normalised to *Gapdh* (F ACCTGCCAAGTATGATGACATCA and R GGTCCTCAGTGTAGCCCAA GAT) mRNA by calculating $2^{-\Delta Ct}$.

*RNA sequencing.* RNA yield, quality and integrity were evaluated using the RNA 6000 Pico kit on an Agilent 2100 Bioanalyser (Agilent Technologies, USA). Only samples with a RNA Integrity Number (RIN) above 8.0 were included in the study. cDNA amplification from total RNA was performed using the SMART-seq® v4 Ultra® Low Input RNA Kit (Clontech, USA). Paired-end sequencing libraries were prepared from the amplified cDNA according to the Nextera® XT DNA library prep protocol (43 bp reads, approx. 23 million reads/sample), and sequenced using an Illumina NextSeq 500 (Illumina, USA).

***RNA sequencing analysis.*** Expression of transcript counts was quantified using *kallisto* (4) software with a GRCh38 transcript model. The data were imported to the R statistical environment and summarized at the gene level (that is transcript counts summed per gene) using *tximport* (5). Statistical transformations for visualisation (for example, the rlog) and preprocessing and then analyzes of differential expression were performed using *DESeq2* (6). Multiple testing adjustments of differential expression utilised the Benjamini–Hochberg (False Discovery Rate) and independent hypothesis weighting (IHW) adjustments (7).

*GeneSet Enrichment Analysis.* We used GeneSet Enrichment Analysis (GSEA) (8) together with the MSigDB pathways database (9). Briefly the statistical output from DESeq2 analysis was used to rank the differential expression of all genes from most upregulated to most downregulated. Mouse genes were then converted to the nearest human homologue and then the ranking used as input for the GSEA tool and the MSigDB 'Hallmark' GeneSets - a curated set of well defined biological processes.

*Gene expression analysis for heat maps.* FASTQ Toolkit, version 1.0.0 (BaseSpace, Illumina), was used for adapter trimming of the reads. Alignment and mapping of all libraries were performed using TopHat Alignment, version 2.0.0, and Cufflinks Assembly & DE, version 2.0.0 (BaseSpace, Illumina, USA). The dataset was filtered to remove genes where two or more samples in each group had an FPKM count of <1. Gene expression levels were calculated using the Cufflinks Assembly & DE, version 2.0.0 (BaseSpace, Illumina, USA), employing a geometric library normalisation method and a fragment bias correction algorithm. Differential gene expression was calculated using a threshold of fold change > 2 and false discovery rate (FDR) < 0.05. Gene expression graphs are shown as relative FPKM counts were normalized to the maximum.

*Overrepresentation analysis.* The Web-based Gene Set Analysis Toolkit (WebGestalt), a suite of tools for functional enrichment analysis, was used to identify overrepresented gene ontology (GO) annotation categories and translate gene lists into functional profiles. Enrichment of GO terms and associated p-values were calculated based on hypergeometric distribution statistics, adjusting the false discovery rate using the Benjamini-Hochberg procedure.

**In vitro T cell assays.** LC were sorted from epidermal single cell suspensions and pulsed with 0.5μM SIINFEKL peptide in T cell medium (RPMI (RPMI 1640, Lonza, Switzerland), 5% heat-inactivated FBS (Life Technologies, USA), 1% L-glutamine (2 mM; Life Technologies, USA), 1% Pen-strep (100 U/ml Life Technologies, USA) and 50μM β-ME (Sigma-Aldrich, UK)) for 30 min at 37°C. Splenic OT-I cells, were directly isolated using the Miltneyi MACS system (OctoMACS Separator, MS columns, CD8a [Ly-2] MicroBeads; Miltenyi Biotec), and labelled with 5μM cell trace violet (Molecular Probes) according to the manufacturer's instructions. $5 \times 10^4$ T cells were co-cultured with 100 LC for 65 hours before analysis by flow cytometry. Cells were gated on live, single $CD45.1^+CD8^+$ T cells.

**Culture of $EpCAM^+$ cells.** Coverslips were pre-coated with 2% (3-Aminopropyl)triethoxysilane (Sigma-Aldrich, UK) in acetone and placed in the wells of 6-well tissue culture plates. These were covered with $5 \times 10^4$ primary keratinocytes (live, $CD45^{neg}$) sorted from the epidermis of untreated female C57BL/6 mice. Approximately 4 hours later, purified $EpCAM^+$ ($Langerin^{GFP}EpCAM^+$

CD24$^{low}$) cells were sorted from BMT+ T cell recipients and added to the wells with 5ng/ml TGFβ. Cells stained and imaged by confocal microscopy using the 40X objective 36 hours later.

**Mathematical modelling.** As described in the text we modelled the flows, and dynamics within, three populations – CD11b$^{high}$ cells, EpCAM+ intermediates, and mature LC. We refer to these populations as $P_1$, $P_2$ and $P_3$ respectively. We considered two developmental pathways – linear ($P_1 \to P_2 \to P_3$) and branched ($P_1 \to P_2$, $P_1 \to P_3$). To describe these, we fitted empirical descriptor functions to the timecourses of numbers of Ki67$^{hi}$ and Ki67$^{lo}$ CD11b$^{high}$ cells, which we refer to $P_1^+(t)$ and $P_2^-(t)$ respectively. These were both of the form

$$\log_{10}(f(t)) = c + \frac{at}{(b+t)^2}$$

Then, for the Ki67$^{hi}$ and Ki67$^{lo}$ EpCAM+ cells (the $P_2$ population), the simplest, general model was

$$\frac{dP_2^+}{dt} = \mu_1 P_1^+ + \rho_2(2P_2^- + P_2^+) - (\beta + \delta_2)P_2^+ \tag{1}$$

$$\frac{dP_2^-}{dt} = \mu_1 P_1^- - \rho_2 P_2^- + \beta P_2^+ - \delta_2 P_2^- \tag{2}$$

where we use the superscripts $+/-$ to denote Ki67$^{hi}$ and Ki67$^{lo}$ cells respectively. Here $\rho_2$ is the rate of cell division; $\beta$ is the mean lifetime of Ki67 expression post-division, which we set at 3.5d (Gossel *et al.*, eLife (2017)); $\mu_1$ is the per capita rate of differentiation from $P_1$ to $P_2$; and $\delta_2$ is the rate at which $P_2$ cells are lost through death or onward differentiation, combined.

We fitted the above equations simultaneously to the total numbers of cells in $P_2 = P_2^+ + P_2^-$, and the Ki67 proportion $P_2 = P_2^+/(P_2^- + P_2^+)$, log- and logit-transformed respectively to normalise residuals. This fitting strategy, detailed in Hogan, Gossel *et al.*, PNAS (2015), maximises the joint log likelihood of the datasets. Fitting on counts and Ki67 fractions rather than the timecourses of Ki67$^{hi}$ and Ki67$^{lo}$ cells assures independence of errors. We found no evidence for cell division within $P_2$ ($\rho_2 \simeq 0$), giving the simpler model

$$\frac{dP_2^+}{dt} = \mu_1 P_1^+ - (\beta + \delta_2)P_2^+ \tag{3}$$

$$\frac{dP_2^-}{dt} = \mu_1 P_1^- + \beta P_2^+ - \delta_2 P_2^-. \tag{4}$$

We then modelled $P_3$ as follows, assuming it is fed from a precursor population $X(t)$, which could be $P_1$ (the branched pathway) or $P_2$ (the linear pathway);

$$\frac{dP_3^+}{dt} = \mu X^+(t) + \frac{\rho_3}{1 + P_3(t)/\lambda}(2P_3^- + P_3^+) - (\beta + \delta_3)P_3^+ \tag{5}$$

$$\frac{dP_3^-}{dt} = \mu X^-(t) - \frac{\rho_3}{1 + P_3(t)/\lambda}P_3^- + \beta P_3^+ - \delta_3 P_3^- \tag{6}$$

where $P_3 = P_3^- + P_3^+$. The term proportional to $\rho_3$ is a density-dependent division rate that decreases with the size of the mature LC pool $P_3$ and is half-maximum when $P_3 = \lambda$. Mature

**Supplemental references.**
